# Advancing Fair and Explainable Machine Learning for Neuroimaging Dementia Pattern Classification in Multi-Ethnic Populations

**DOI:** 10.1101/2025.06.09.658375

**Authors:** Ngoc-Huynh Ho, Sokratis Charisis, Nicolas Honnorat, Sachintha Ransara Brandigampala, Di Wang, Susan R. Heckbert, Peter T. Fox, David Martinez, David H. Wang, Timothy M. Hughes, Derek B. Archer, Timothy J. Hohman, Sudha Seshadri, Christos Davatzikos, Mohamad Habes

## Abstract

Dementia, a degenerative disease affecting millions globally, is projected to triple by 2050. Early and precise diagnosis is essential for effective treatment and improved quality of life. However, current diagnostic approaches frequently demonstrate inconsistent precision and impartiality, particularly among diverse cultural groups. This study investigates performance discrepancies in dementia classification among White American, African American, and Hispanic populations. We reveal significant cross-group bias, particularly when models trained on one group are tested on another. To address this, we introduce a novel combination of few-shot learning and domain alignment to improve model adaptability across underrepresented populations. Our results show that these techniques substantially reduce inter-group performance gaps, especially between White American and Hispanic cohorts. This finding highlights the crucial need for fairness-aware strategies and the inclusion of diverse populations in training data to ensure accurate and equitable dementia diagnoses.

## Introduction

Dementia is a clinical condition characterized by a gradual decline in cognitive function that affects memory, reasoning, communication, and the ability to perform daily tasks. This decline is typically the result of brain cell damage from conditions such as Alzheimer’s disease (AD) ^1–5^. Early diagnosis can play a crucial role in customizing treatment plans and managing the disease more effectively ^6–8^. In recent years, artificial intelligence (AI) has shown promising potential in improving the accuracy of dementia diagnoses by applying advanced machine learning (ML) techniques to medical imaging data, including magnetic resonance imaging (MRI). These models are capable of detecting subtle, often subclinical, structural brain changes, enabling earlier and more reliable diagnoses than traditional clinical approaches ^9–12^.

Nevertheless, AI-driven diagnostic systems are subject to inconsistent outcomes, particularly when applied across various cultural contexts ^13–18^. These inconsistencies refer to systematic inaccuracies in classifying or representing specific demographic groups, often defined by factors such as ancestry, age, gender, culture, socioeconomic, and education, within diagnostic algorithms ^19–21^. These inaccuracies may result in either underdiagnosis or overdiagnosis ^22, 23^, ultimately leading to suboptimal clinical outcomes and amplifying existing healthcare discrepancies. In the context of dementia, anatomical differences among groups and the limited representation of certain demographic segments in medical datasets can contribute to imbalanced diagnostic performance. Mayeda et al. ^24^ reported significant variations in dementia incidence across six population groups over 14 years. Similarly, Gilsanz et al. ^25^ explored dementia rates in individuals aged 90 and above in a broadly representative cohort, offering insight into how risk varies across diverse backgrounds.

Emerging research has highlighted significant variations in Alzheimer’s disease diagnoses between population groups. For instance, while the presence of the *APOE*-*ε*4 allele is a recognized risk factor among individuals of White descent ^15, 26, 27^, studies have found that African American and Hispanic populations exhibit elevated Alzheimer’s rates regardless of APOE genotype ^28, 29^. These studies suggest potential variations in genetic or environmental risk factors among these populations. As previously described^27^, differences in brain morphology, economic conditions, and access to care are influential in shaping these outcomes. AI-based diagnosis tools can sometimes amplify these discrepancies. For example, ML models predominantly trained on data from White American populations often exhibit reduced accuracy when applied to other demographic groups^30^. The study conducted by Obermeyer et al.^31^ highlighted how race-neutral healthcare algorithms can unintentionally perpetuate diagnostic imbalances. When predicting colorectal cancer, Khor et al. ^32^ measured higher false-positive rates for Hispanic and Asian patients compared to non-Hispanic White patients in statistical models that did not take ethnicity into account.

In this study, we explore diagnostic inconsistencies in dementia classification by analyzing MRI-derived features from White American, African American, and Hispanic populations. ML models were applied to data from 3, 127 participants to identify discrepancies in diagnostic outcomes. To address these diagnostic issues, we employed data harmonization to standardize MRI inputs, correlation removal techniques to reduce the influence of nuisance characteristics on predictions, and domain adaptation strategies, including kernel mean matching, semi-supervised learning, and few-shot learning, to improve generalization. These interventions aimed to maintain diagnostic accuracy while enhancing consistent and balanced diagnostic outcomes across groups. Lastly, we used advanced feature visualization methods to identify the brain regions associated with the ethnic group discrepancies. This work highlights the importance of heterogeneity in model training and discrepancy mitigation strategies in developing fair, representative, and reliable AI applications for dementia diagnosis and care across diverse population groups.

## Methods

The proposed framework consists of three main components: (a) an end-to-end pipeline including preprocessing steps such as reorientation, bias correction, and brain region extraction using Multi-Atlas Region Segmentation using Ensembles (MUSE) ^33^; (b) a model training relying on Extreme Gradient Boosting (XGBoost) ^34^ combined with a discrepancy mitigation technique; and (c) a comprehensive evaluation of model fairness and interpretability, carried out using data across multiple AD research centers, which outlines two cross-validation strategies, training on all populations versus training on a single group, to assess model generalizability and classification discrepancies. The detailed training procedures are provided in the Experimental Settings section (Supplementary). We summarized this proposed framework for fair dementia classification in Figure 1.

**Figure 1:**
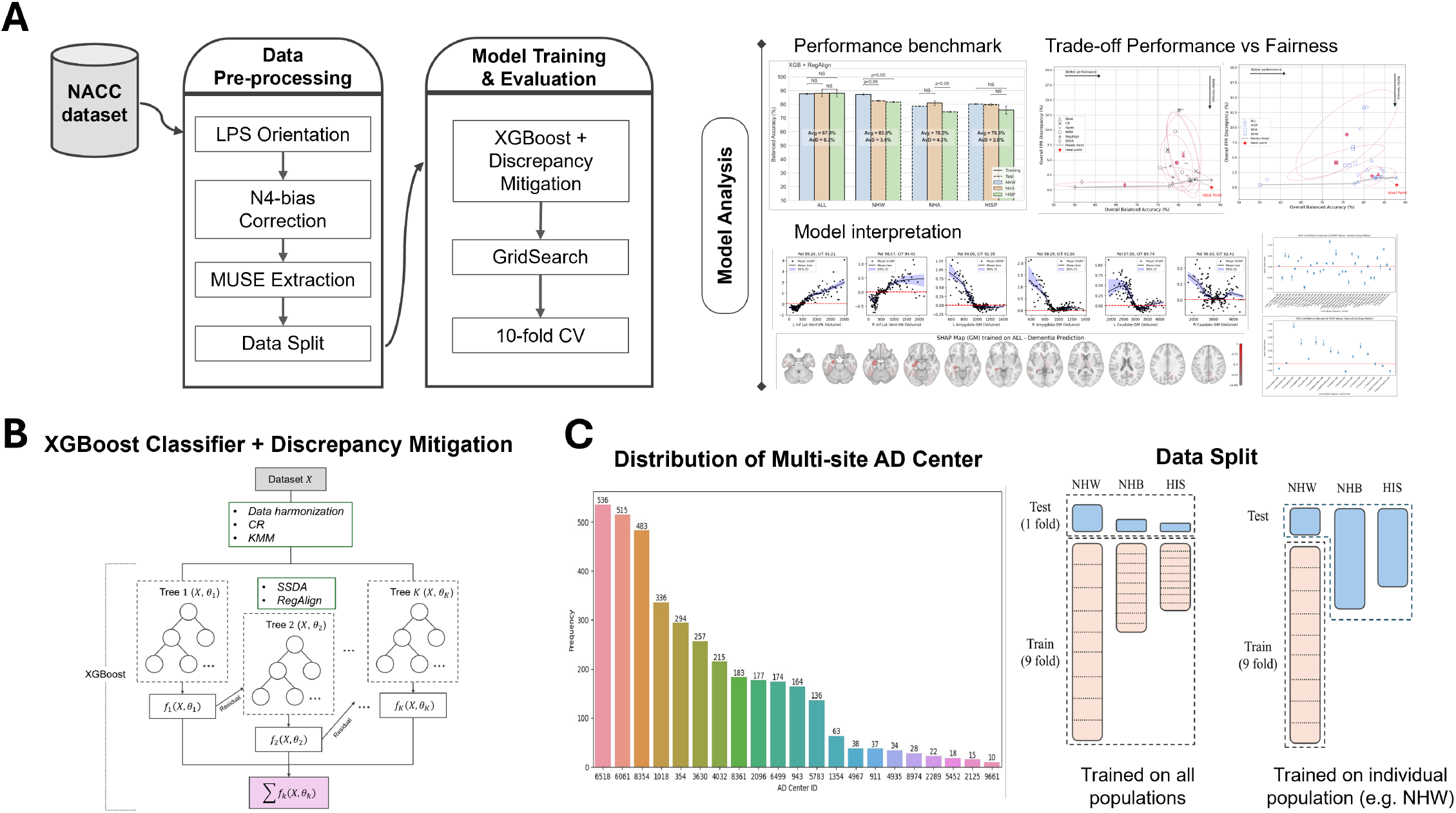
Overview of the proposed machine learning framework for fair dementia classification. **A** – The study uses multi-ethnic MRI data from the NACC dataset. The pipeline integrates preprocessing, domain-aware training with a customized loss (RegAlign), and interpretable evaluation using SHAP-based brain region importance. Evaluation includes performance metrics across racial/ethnic subgroups, fairness assessments via Pareto analysis, and visualization of feature contributions. **B** – Visualization of the XGBoost classifier combined with the customized loss function, RegAlign. **C** – Multi-site data distribution and the strategy for training/test splitting.

### Study Populations

Participants were obtained from the NACC database (http://www.alz.washingfaton.edu/). The data included patients assessed at Alzheimer’s Disease Centers (ADCs) between January 2005 and June 2012. The NACC recruitment and data-gathering process, which spans multiple ADC sites, has been previously described in ^35^. In this study, participants were collected from 21 ADCs, emphasizing the dataset’s center multiplicity, as shown in Figure 1 (Panel C). The study consisted of 2, 007 participants with “normal cognition” (mean age = 69.03) and 1, 120 participants with “dementia” (mean age = 73.06) from the Uniform Data Set (UDS). To investigate potential disparities in dementia classification, we filtered the cohort to include participants from three self-reported racial/ethnic groups: Non-Hispanic White American (NHW), Non-Hispanic African American (NHA), and Hispanic (HISP). Table 1 provides a comprehensive summary of participant demographics, including age, sex, education, and Mini-Mental State Examination (MMSE) scores.

**Table 1:** Demographic and clinical characteristics by race/ethnicity.

| Variable |  | NHW | NHA | HISP | p-value |
| --- | --- | --- | --- | --- | --- |
| Study participants (n) |  | **2,530** | **337** | **249** | – |
| Sex (M\|F) | | 1,050 \| 1,480 | 79 \| 258 | 80 \| 169 | $p < 0.001^{\dagger}$ |
| Age (std), years | NC | 69.02 (12.68) | 71.66 (7.8) | 70.13 (10.6) | $p < 0.01^{\ddagger}$ |
| | Dem | 73.06 (11) | 75.57 (9.57) | 73.52 (10.9) | $p = 0.107^{\ddagger}$ |
| Education (std), years | NC | 16.34 (2.52) | 15.22 (2.84) | 11.80 (4.87) | $p < 0.001^{\ddagger}$ |
| | Dem | 15.40 (2.95) | 14.04 (2.93) | 11.89 (5.01) | $p < 0.001^{\ddagger}$ |
| MMSE (std) | NC | 29.07 (1.27) | 28.11 (1.58) | 27.77 (1.95) | $p < 0.001^{\ddagger}$ |
| | Dem | 20.93 (5.56) | 18.00 (6.61) | 18.47 (6.49) | $p < 0.001^{\ddagger}$ |
| Dementia ratio (%) |  | 35.85% | 27.3% | 45.78% | – |

NC : Normal Cognition • Dem : Dementia • std: standard deviation
NHW : Non-Hispanic White • NHA = Non-Hispanic African • HISP = Hispanic
<sup>†</sup> p-values were calculated using the $\chi^2$ test for categorical variables.
<sup>‡</sup> p-values were calculated using one-way ANOVA for continuous variables.

### Image Processing

The study utilized raw T1-weighted MRI data in DICOM format from the NACC database to facilitate an in-depth investigation of brain regions of interest (ROIs). We first reoriented the images according to the LPS (Left-Posterior-Superior) convention to standardize their spatial orientation. N4-bias field correction ^36^ was employed to reduce intensity inhomogeneity across the images, leading to more precise post-processing. Next, we performed skull stripping and brain segmentation using Multi-Atlas Region Segmentation using Ensembles (MUSE) ^33, 37^, which combines multiple atlas-based segmentations to improve robustness and accuracy. MUSE generated 144 region-of-interest (ROI) features, providing detailed and reproducible anatomical parcellations relevant to neurological research. These features were subsequently used for Alzheimer’s disease classification, supporting cross-group comparisons of brain atrophy patterns among different racial and ethnic populations ^38^.

### ML Classifier and Discrepancy Mitigation Methods

In this study, we used XGBoost (XGB), a gradient-boosting technique recognized for its ability to manage imbalanced datasets ^34, 39, 40^, as a baseline classifier for dementia classification. We trained the model to distinguish between normal cognition (NC) and dementia (Dem) classes utilizing MUSE-based brain ROI extraction from MRI scans. These features capture structural variations across 144 anatomical regions and serve as a biologically informed input representation for classification. XGB was selected not only for its strong generalization performance but also for its ability to model non-linear interactions between brain regions. Moreover, XGB allows for the derivation of feature importance rankings, enabling us to identify and interpret the brain alterations most strongly associated with dementia.

#### Conventional Approaches for Discrepancy Mitigation

To address the diagnosis discrepancy in dementia classification among racial and ethnic groups, we implemented five bias mitigation strategies, including data harmonization, correlation remover, kernel mean matching, semi-supervised domain adaptation, and the proposed RegAlign. Data harmonization was performed via Combat-GAM ^41–43^, a technique that compensates for site-specific and demographic factors (such as age, gender, race/ethnicity, and diagnosis) to mitigate batch effects and maintain uniformity across multi-site MRI data ^44, 45^. This technique harmonized data distributions among racial and ethnic groups, thus reducing data heterogeneity.

Correlation Remover (CR) ^46–48^ algorithm uses a linear transformation to remove correlations between sensitive features (e.g., race/ethnicity) and non-sensitive features (e.g., MRI-derived brain regions). CR ensures, therefore, that the sensitive features do not influence the non-sensitive ones. Formally, let the dataset **X** consist of two subsets: sensitive features **S** and non-sensitive features **Z**. Let *m*_*s*_ and *m*_*z*_ denote the number of sensitive and non-sensitive features, respectively. The mean of the sensitive features is defined as 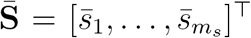, where 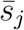 is the mean of the *j*-th sensitive feature. For each non-sensitive feature **z**_*j*_ ∈ ℝ^*n*^, where *j* = 1, …, *m*_*z*_, an optimal weight vector 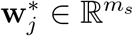 is calculated by minimizing the least squares objective:

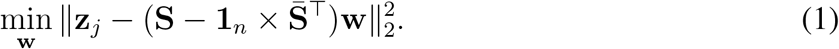

Here, **1**_*n*_ is a vector of ones in ℝ^*n*^. In this formulation, 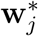 represents the solution to a linear regression problem where the non-sensitive feature **z**_*j*_ is projected onto the sensitive features, which are mean-centered. By solving this problem for all non-sensitive features, the weight matrix 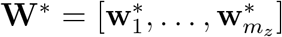 is derived. Using this weight matrix, the adjusted non-sensitive features **Z**^*^is computed as:

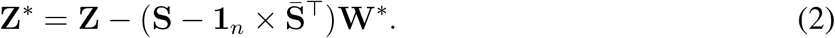

Finally, the sensitive features **S** are removed from the dataset **X**, and **Z**^*^ replaces the original non-sensitive features. A hyperparameter *α* (where *α* = 0.05) is introduced to allow flexibility in the degree of filtering applied, defined as:

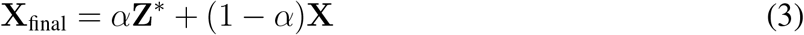

Besides, the domain adaptation technique ^49^ was applied using Kernel Mean Matching (KMM) ^50, 51^ to further mitigate bias. KMM is designed to address the problem of distribution mismatch between source and target domains in machine learning tasks. It reweights the source domain samples to more closely align with the target domain’s distribution in a reproducing kernel Hilbert space (RKHS) ^52^, ensuring that the model performs well across different racial and ethnic groups. Let *ϕ*(*x*) be a feature mapping to a RKHS ℋ. The objective is to find weights *β* that minimize:

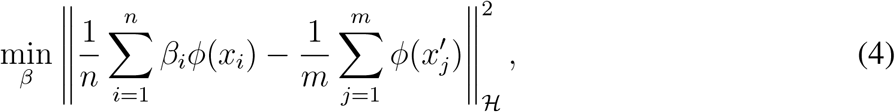

while ensuring the following conditions are met:

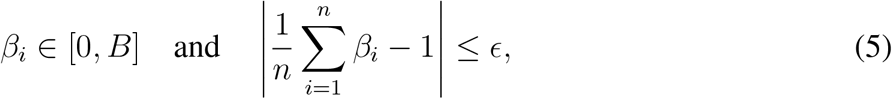

Where *x*_*i*_ and 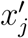 are source and target domain samples, *n* and *m* are the number of source and target samples, *B* is the upper bound on the weights and was set to 512, and *ϵ* controls the discrepancy between the mean weights and 1, and is computed as 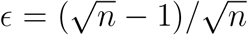. The default radial basis function (RBF) kernel is utilized. The estimated sample weights are subsequently normalized to ensure that their total equals 1, thereby preserving the relative differences between samples while maintaining a stable and interpretable learning process. Then, the final sample weights are served in the ML classifier to modify the impact of each training sample throughout the model’s learning process. TThe loss function is adjusted by assigning weights that reflect the importance of each sample. For instance, samples with higher weights significantly impact the overall loss, drawing the model’s focus, while lower-weight samples have a reduced influence.

The semi-supervised domain adaptation (SSDA) method is inspired by entropy minimization ^53^, which encourages confident (i.e., low-entropy) predictions on the unlabeled target domain. Given *n* labeled source samples (*x*_*i*_, *y*_*i*_) and *m* unlabeled target samples *x*_*j*_, the total loss includes:

##### (1) Supervised loss on source domain

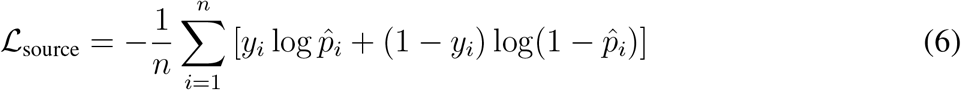

##### (2) Entropy minimization on target

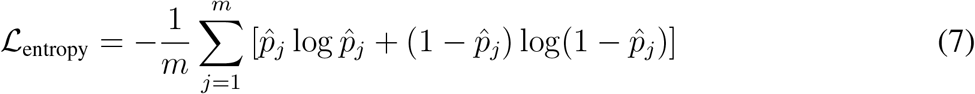

##### (3) Class-conditional alignment using pseudo-labels

Binary pseudo-label 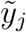 is generated from the target predictions (e.g., using K-means clustering), and compute the predicted mean *µ*^(*c*)^ for each class *c* ∈ {0, 1} in both source and target:

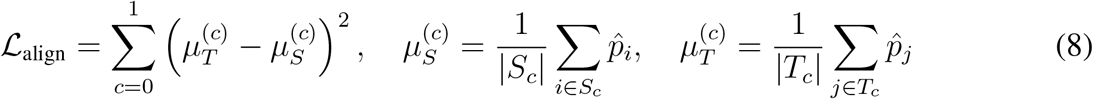

where the sets *S*_*c*_ and *T*_*c*_ represent the index sets of class-*c* samples in the source and target domains, respectively.

The total loss is aligned by control hyperparameters *τ* and *λ* as follows:

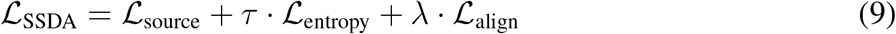

#### Proposed RegAlign

In typical domain adaptation settings, training on a source population using standard loss functions (e.g., cross-entropy or focal loss) often leads to reduced performance on underrepresented target populations, particularly when only a few labeled samples from the target domain are available. These issues are exacerbated when the source domain (e.g., African American or Hispanic cohorts) exhibits class imbalance, which causes models to overfit to the majority class (often normal cognition, NC) and underperform on the minority class (dementia). To address these challenges, we propose a custom loss function, termed RegAlign, which integrates three key components: (1) focal weighting to emphasize hard-to-classify samples within the source domain, (2) class-weighted binary cross-entropy to minimize false negatives in the few-shot target domain, and (3) a regularized alignment term to match class-conditional prediction distributions between source and target domains. By combining these elements, RegAlign performs few-shot domain adaptation more effectively and ensures balanced optimization across source and target datasets.

##### (1) Focal-weighted BCE on source

To improve robustness against class imbalance (common in minoritized groups), we use focal weighting on the source loss:

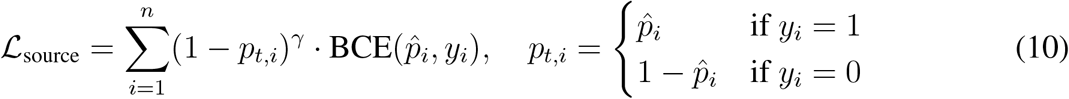

##### (2) Class-weighted BCE on few-shot target

To reduce false negatives, we up weight positive class examples using a scalar *α* > 1:

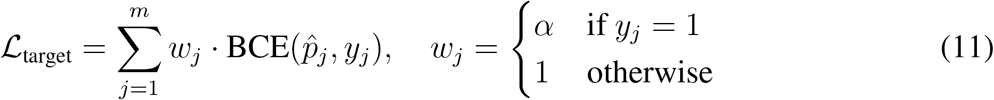

##### (3) Regularized class-conditional alignment

We align mean predictions of class-0 and class-1 between source and target domains using Eq. 8. This term is scaled by a regularization coefficient *λ* ∈ (0, 1) to prevent overfitting to few-shot target data.

##### Final loss

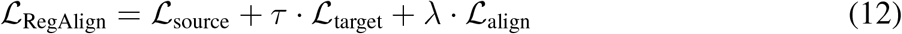

### Model Interpretation via SHAP

To interpret model decisions and identify critical brain regions influencing dementia classification, we employed SHapley Additive exPlanations (SHAP) ^54^. We calculated SHAP values for a baseline XGBoost classifier under the four distinct training configurations described in detail in the first supplementary, Experimental Settings section. The resulting attribution maps highlight how each brain region contributes to the model’s decision in classifying individuals as having dementia. Red regions on these maps indicate features pushing the prediction toward a specific class, while the color intensity reflects the magnitude of the SHAP values.

To efficiently compute these SHAP values, particularly on GPU-enabled systems, we implemented a custom PyTorch-based routine that approximates Tree SHAP using perturbation-based differences^55, 56^. This method is conceptually similar to conventional SHAP estimation, where a feature’s contribution is determined by perturbing its value and measuring the change in prediction. Specifically, the SHAP value for each feature is estimated by comparing model predictions before and after replacing that feature with its average across a reference background dataset *G* ∈ ℝ^*m*×*d*^, typically the training set. Let *f* : ℝ^*d*^ → ℝ be the model’s score prediction function, and let *X* ∈ ℝ^*n*×*d*^ be a batch of input instances. For each input *x* ∈ X ⊂ ℝ^*n*×*d*^ and feature index *j*, we generate a perturbed copy *x*^′^ by replacing *x*_*j*_ with the average value from *G*. The SHAP value *ψ*_*x,j*_ for feature *j* in instance *x* is then estimated as the prediction difference:

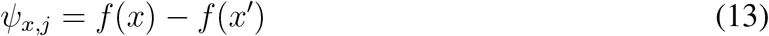

#### Algorithm

SHAP Computation

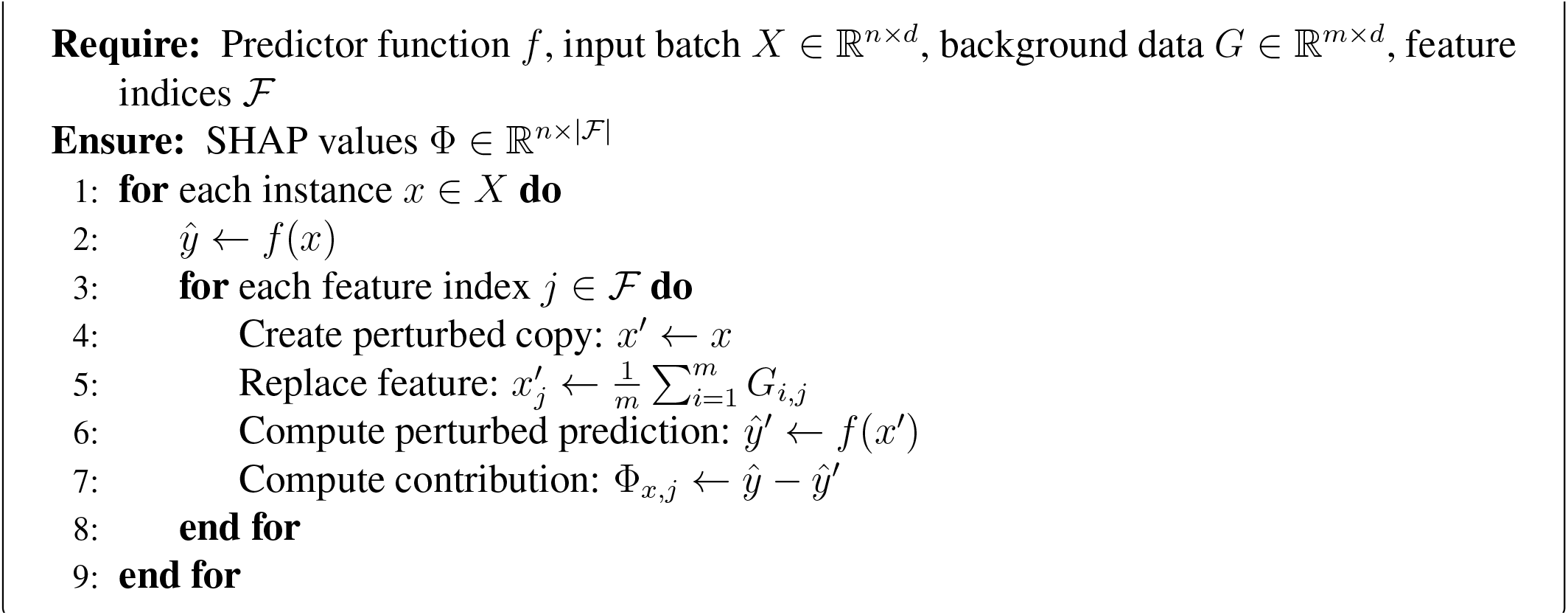

where and Ψ ∈ ℝ^*n*×|*F* |^ stores SHAP values for all instances and selected features. This method enables localized SHAP estimation for GPU-enabled XGBoost models, facilitating the identification of significant brain regions that influence the classifier’s outputs. To capture the uncertainty in SHAP estimates, we applied a non-parametric bootstrap method based on residual resampling^57, 58^. Specifically, the residuals are defined as:

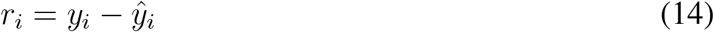

where *y*_*i*_ is the true label and 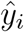 is the model’s prediction for instance *i*. These residuals represent the model’s unexplained errors. To generate a bootstrapped target vector 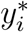, we resample the residuals *r*_*i*_ with replacement and add them back to the original predicted values:

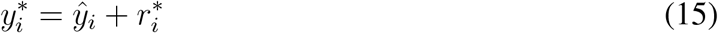

where 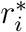 is a resampled residual. Then, a new model is retrained on (*X, y*^*^) using the same hyperparameters, and SHAP values are recomputed. This procedure is repeated 5,000 times to generate a bootstrap distribution of SHAP values and estimate feature-level uncertainty.

In addition, we generated a preliminary meta-analysis map by reprocessing a set of voxel-based morphometry (VBM) studies collected from a previous meta-analysis ^38^. We created a brain map by applying the activation likelihood estimation (ALE) method, using the GingerALE software (version 3.0.2) ^59^, to combine the selected VBM studies into a single ALE map that highlights brain regions where atrophy is likely associated with Alzheimer’s disease. The ALE analysis was conducted using a cluster-forming threshold of 0.001 and a cluster-level significance threshold of 0.05, with statistical significance estimated through 1, 000 random permutations, following previous practices ^38^. The continuous ALE map produced by GingerALE, in which non-significant brain areas were assigned null values, was then thresholded at the smallest non-zero value to create a binary map, facilitating comparisons across different training strategies.

### Evaluation metrics

Dementia classification performance was assessed using two fairness measures and balanced accuracy (BA). BA, the average of sensitivity or true positive rate (TPR) and specificity or true negative rate (TNR), captures variances between normally cognitive and demented classes, offering a more comprehensive evaluation than standard accuracy. This metric proved effective in handling the unbalanced datasets observed in this study.

To assess the fairness of the models across three racial and ethnic groups, we examined disparities in the false positive rate (FPR) and false negative rate (FNR) between the groups, similar to previous studies ^18, 21, 21^. FPR has important implications in clinical trials, such as differential access to appropriate care or misallocation of medical resources ^21^. This misallocation not only exposes healthy individuals to potential side effects but also reallocates essential resources away from patients who genuinely need intervention. Similarly, FNR has critical clinical implications, as it can lead to underdiagnosis, delaying access to timely interventions and appropriate care for individuals who may have dementia. In this study, FPR and FNR discrepancies were calculated bu comparing each pair of groups: NHW vs. NHA, NHW vs. HISP, and NHA vs. HISP. The differences from these three comparisons were then averaged to calculate a single metric per group, referred to as the average discrepancy (AvD). Additionally, GridSearchCV is used for hyperparameter tuning with 10-fold cross-validation to achieve optimal model performance across multiple training and testing splits. This hyper-parameter tuning is described in Table S1 (Supplementary).

## Results

### Performance on Dementia Classification

This section evaluates model performance on dementia classification under different training group compositions (shown in Figure 1 and detailed in the Supplementary Materials, Section “Experimental Settings”) and explores methods to mitigate intergroup discrepancies in predictive accuracy and error rates.

#### Potential Discrepancy

In Figure 2, the first three plots illustrate the performance of the XGBoost Classifier (XGB) across four training scenarios: training with all racial and ethnic groups, the NHW group only, the NHA group only, and the HISP group only. The term “Training” refers to performance on the testing subset drawn from the same group as the training data, while “Test” reflects performance on groups different from the training set. When trained on all groups, the XGB model achieved an average BA score of approximately 84.3%. However, when trained on an individual group, its performance declined substantially. Specifically, the BA slightly dropped to 82.7%, 77.3%, and 80% for NHW-, NHA-, and HISP-only training, respectively. Correspondingly, FPR increases ranged from 0.7% to 12.9%, with the HISP-only model performing worst, and FNR increases ranged from 2.4% to 11.9%, with the NHA-only model performing worst. In addition, training with individual populations resulted in increased gaps across groups. Specifically, the NHA-only model exhibited the highest AvD for both BA (9.4%) and FNR (12.3%) scores. Conversely, the HISP-only model demonstrated the largest AvD for the FPR metric, reaching 13.2%.

**Figure 2:**
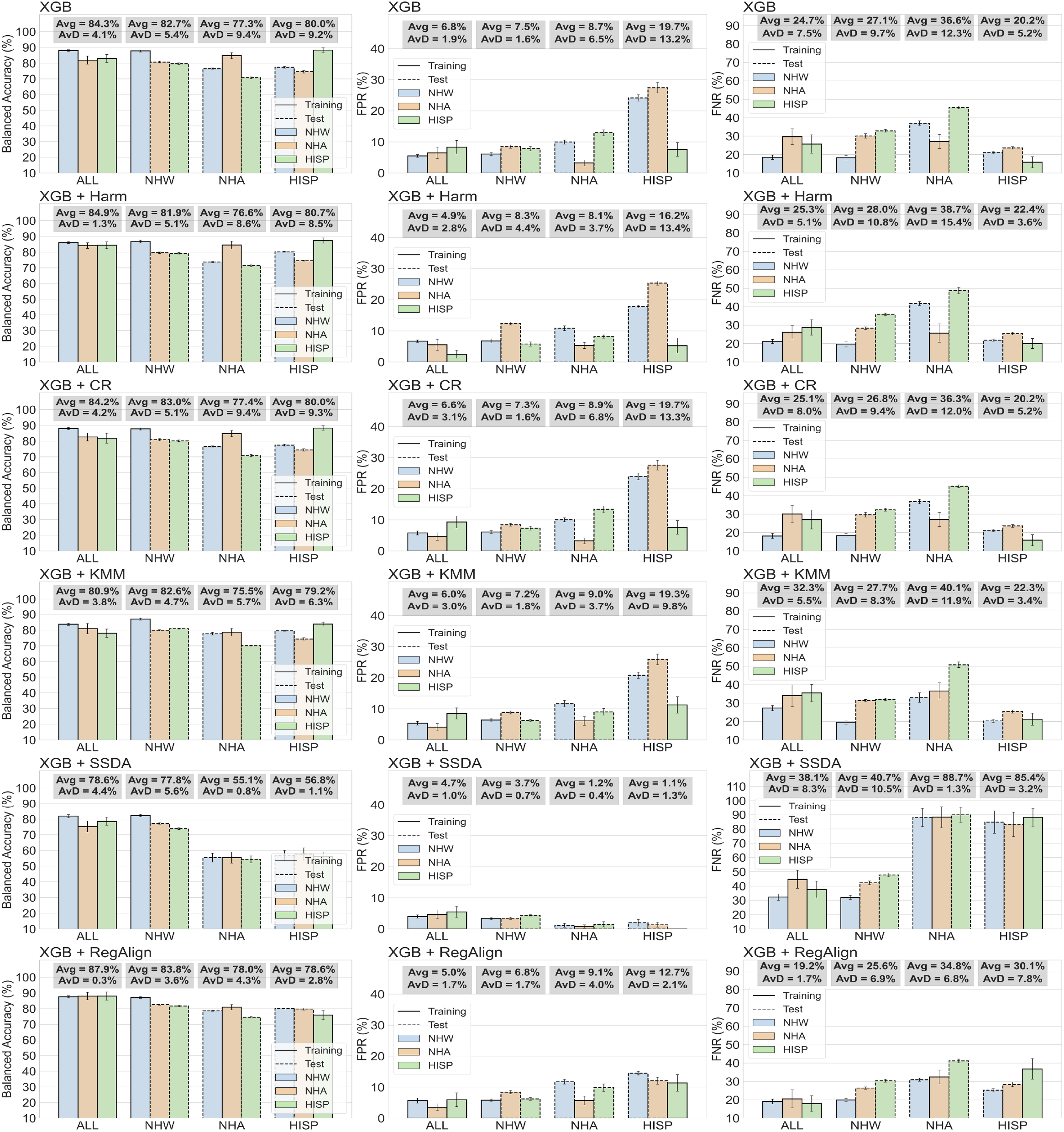
Comparison of baseline classifier and five discrepancy mitigation methods across three evaluation metrics: Balanced Accuracy, false positive rate (FPR), and false negative rate (FNR). The four scenarios include training on all racial/ethnic groups (ALL), or training exclusively on Non-Hispanic White (NHW), Non-Hispanic African (NHA), and Hispanic (HISP) American groups. Where XGB : XgBoost classifier · Harm: data harmonization · CR : correlation removal · KMM kernal mean matching · SSDA : semi-supervised domain adptation · RegAlign : regularized domain alignment · Avg : average value of metric· AvD : average discrepancy. 21

#### Discrepancy Mitigation

Next, we evaluated five discrepancy mitigation strategies as described in the Methods section, aiming to improve both average performance and reduce inter-group differences. Among the evaluated methods, the proposed RegAlign approach achieved substantial improvements in both balanced accuracy (BA) and false negative rate (FNR) when trained on all groups. Specifically, it reached a BA of 87.9% with a minimal AvD of 0.3% and an FNR of 19.2% with an AvD of only 1.7%, indicating both high performance and low inter-group variability. RegAlign also demonstrated a substantial trade-off between accuracy and fairness when trained on individual groups, particularly in BA and FPR metrics. For instance, in the HISP-only training scenario, RegAlign achieved a low FPR of 12.7% with an AvD of 2.1%, outperforming other methods such as data harmonization, CR, and KMM and ranking second only to SSDA. However, the SSDA model achieved a lower FPR partly due to overfitting on the positive class, which led to a significantly higher FNR of 85.4%. This imbalanced performance in the NHA- and HISP-only settings is attributed to the limited number of labeled samples, which prevented the SSDA method from effectively learning the true label distribution. As a result, while SSDA minimized FPR, it compromised sensitivity, making RegAlign a more balanced and robust solution under limited data conditions.

### Trade-off Between Accuracy and Fairness

To evaluate the trade-off between classification performance and fairness, we visualize the Pareto front in Figure 3. In this plot, each method and training scenario is summarized by a confidence ellipse around its distribution, and the distance from a centroid to the ideal point (marked in red start) is used as a quantitative proxy for optimality. The left panels display method-wise comparisons, while the right panels compare training scenarios. As shown in Figure 3, the proposed *RegAlign* achieves the most optimal trade-off, exhibiting the shortest Euclidean distance to the ideal point (high accuracy, low discrepancy) across both FPR and FNR, followed the application of the KMM approach. When considering scenario-based trade-off analysis, training on all racial/ethnic populations yields a more ba trade-off compared to training on any single population. These results align with previous findings: diverse and inclusive training data, combined with few-shot domain adaptation using a customized loss function like RegAlign, is crucial for optimizing both fairness and performance in dementia classification.

**Figure 3:**
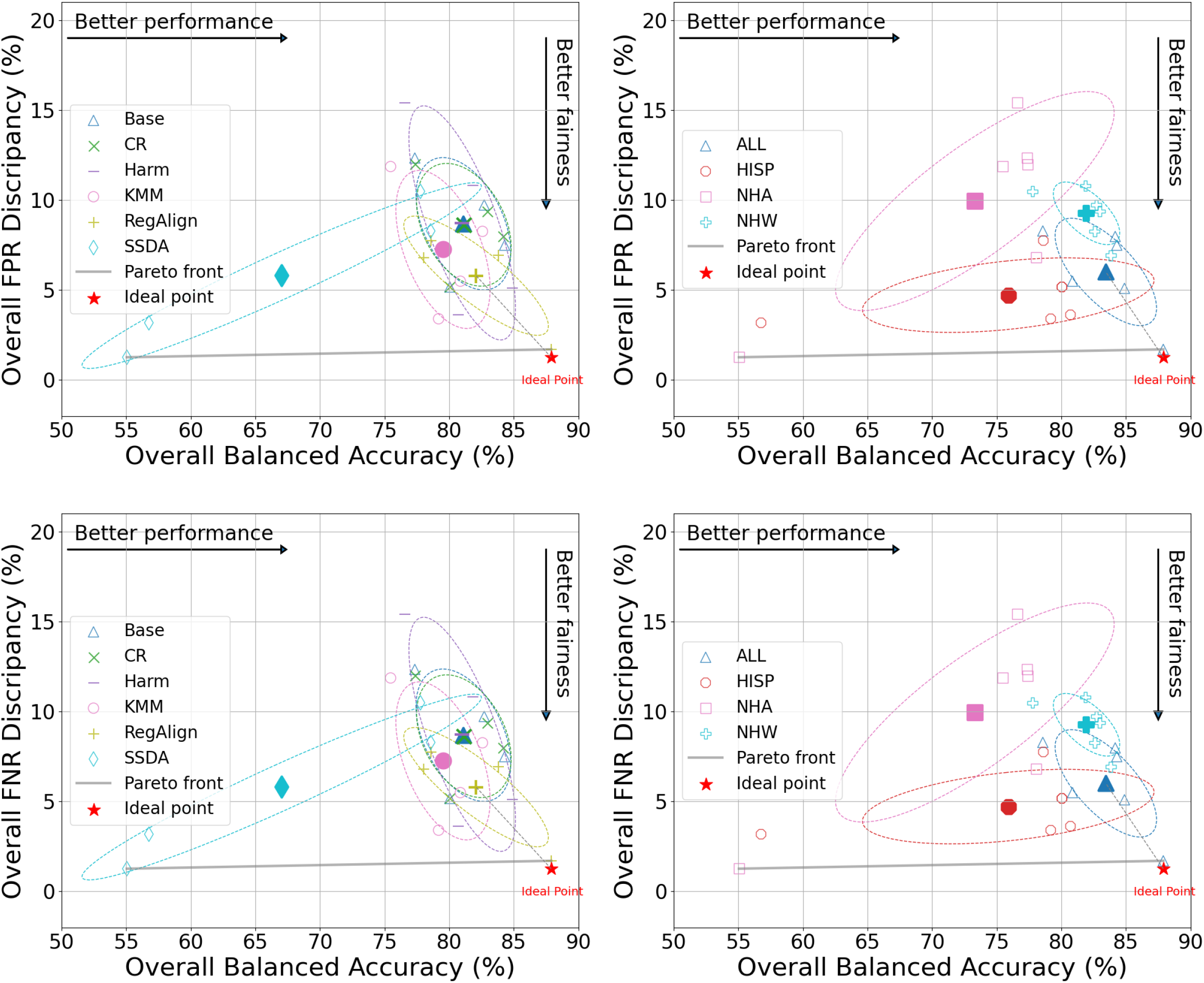
Pareto front optimization for balancing performance and fairness based on FPR (false positive rate) / FNR (false negative rate) discrepancy. Ideal point is defined by the interaction of the highest performance and the lowest fairness. The bigger solid marker denotes the centroid of each group of method or training scenario, the gray dashed line indicates the shortest distance from a centroid to the ideal point, and the ellipse represents the group confidence distribution. Where Harm: data harmonization · CR : correlation removal · KMM kernal mean matching · SSDA : semi-supervised domain adptation · RegAlign : regularized domain alignment.

### Visualization of Brain Region Contributions

In this section, we present the visualization of brain region contributions to the predictive model. By mapping the relative importance of each brain region, we aim to provide an intuitive understanding of how specific anatomical areas influence the classification outcomes. To this end, we computed SHAP values for each brain region feature, following the interpretation framework described in Section *Model Interpretation via SHAP*. The 95% confidence intervals were estimated using empirical percentiles derived from 5, 000 boot-strapping iterations. Only regions whose intervals did not cross zero were considered significant, as shown in Figure S2 (Supplementary).

Next, the ALE meta-analysis identified several brain regions with statistically significant activation likelihood associated with dementia. We found the highest convergence in the left medial temporal lobe, which showed the highest ALE value (0.0556) and *z*-score (8.02), with peak coordinates at (–28, –14, –14) corresponding to the parahippocampal gyrus and hippocampus. Additional significant peaks included the right pallidum (ALE = 0.0264, *z* = 4.74), right supramarginal gyrus (ALE = 0.0245, *z* = 4.49), right amygdala (ALE = 0.0203, *z* = 3.89), and right superior temporal gyrus (ALE = 0.0201, *z* = 3.88).

Figure 4 illustrates the ALE meta-analysis map and the average SHAP maps across four training scenarios: models trained on all populations and those trained exclusively on NHW, NHA, or HISP populations. All images are displayed in MNI space following spatial normalization to ensure anatomical comparability. Each scenario includes paired heatmaps that highlight the contributions of brain regions to predicting dementia. The ALE meta-analysis map is displayed in the upper row as a reference, highlighting consistently reported atrophic regions across VBM studies. When trained on all populations or NHW one, the model predominantly focused on the right hippocampus and parahippocampal gyrus for dementia classification.

**Figure 4:**
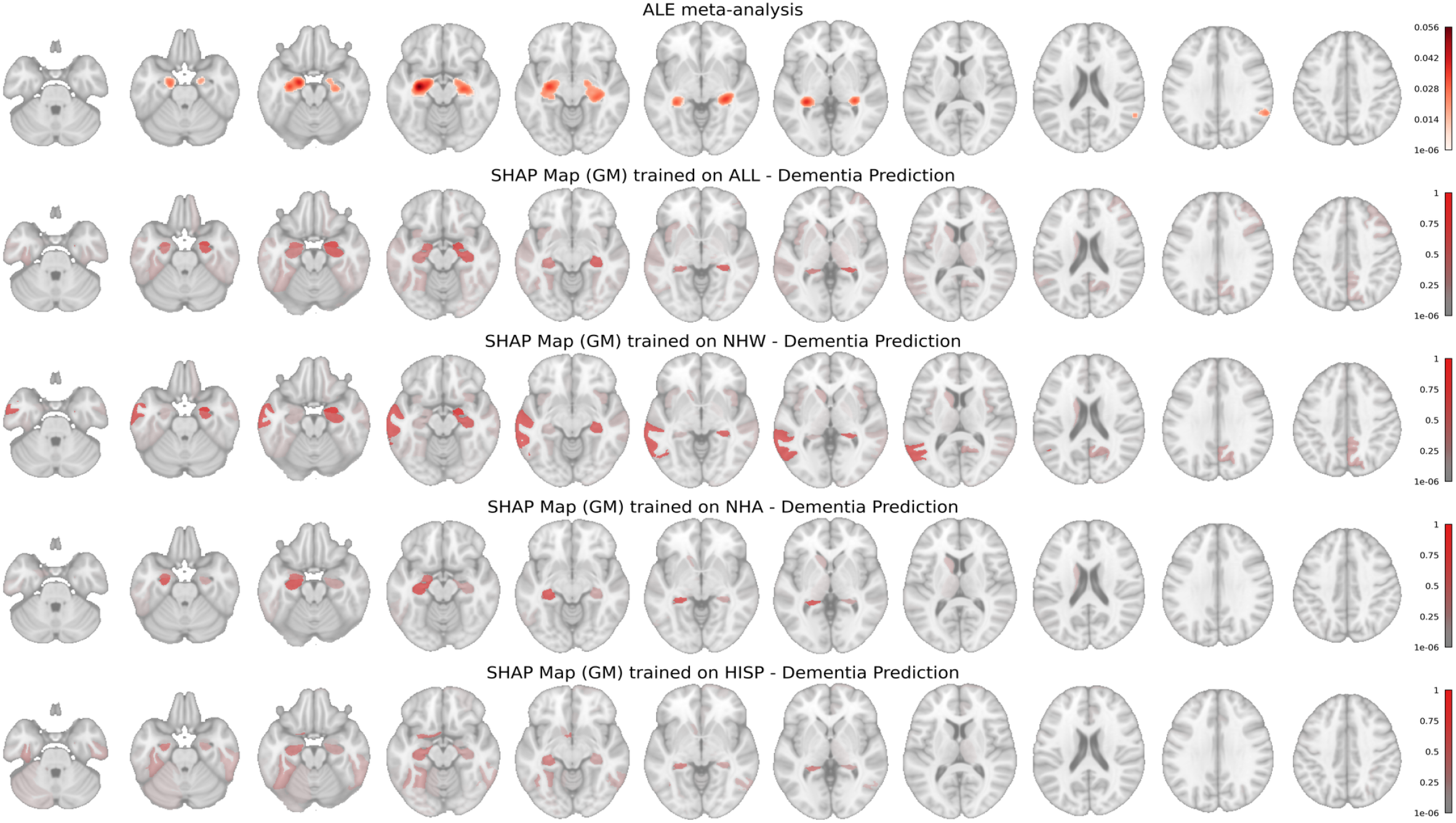
SHAP maps of gray matter region (no ALE-guided) contributions to dementia prediction using the baseline classifier, alongside the ALE template map for reference. Red regions on these maps indicate features pushing the prediction toward a specific class, while the color intensity reflects the magnitude of the SHAP values. Where ALE : activation likelihood estimation · SHAP : SHapley Additive exPlanations · ALL : all racial/ethnic populations · NHW : Non-Hispanic White American · NHA : Non-Hispanic African American · HISP : Hispanic.

In contrast, the model trained on NHA data emphasized the left hippocampus, while the HISP-trained model showed bilateral hippocampal regions. The complete sets of SHAP maps are provided in Figure S5 (Supplementaty). These findings suggest that the model identifies population-specific structural brain changes associated with dementia, potentially leading to biased classifications when trained on one group and tested on another.

Furthermore, Figure 5 presents partial dependence plots of eight dementia-associated brain regions derived from ALE meta-analysis across four population-specific training scenarios: All, NHW, NHA, and HISP. Each plot displays regional brain volume (x-axis) against SHAP values (y-axis), where positive values indicate contributions toward dementia classification and negative values support NC prediction. Across all scenarios, the right and left hippocampus consistently exhibit strong negative SHAP values at lower volumes, aligning with the established role of hippocampal atrophy in dementia diagnosis. Moreover, in the All-trained and NHA-trained models, both hippocampi exhibit steep SHAP transitions from positive to negative values between 3000–3500 mm^3^, suggesting a volume threshold below which the model robustly infers dementia risk. The NHW-trained model shows a similar but slightly weaker distinction, while the HISP-trained model captures this relationship with lower consistency, likely due to sample size limitations.

**Figure 5:**
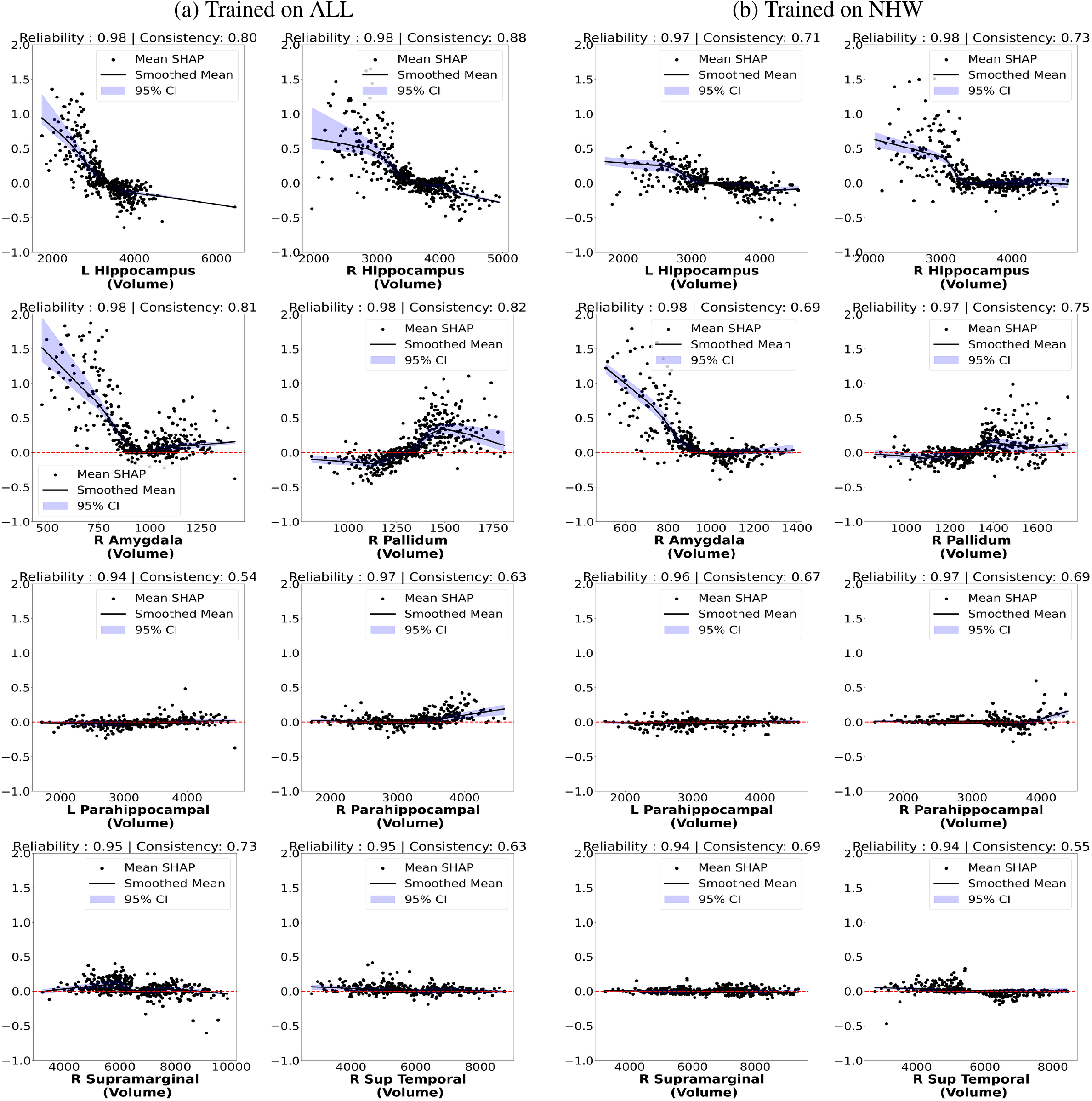
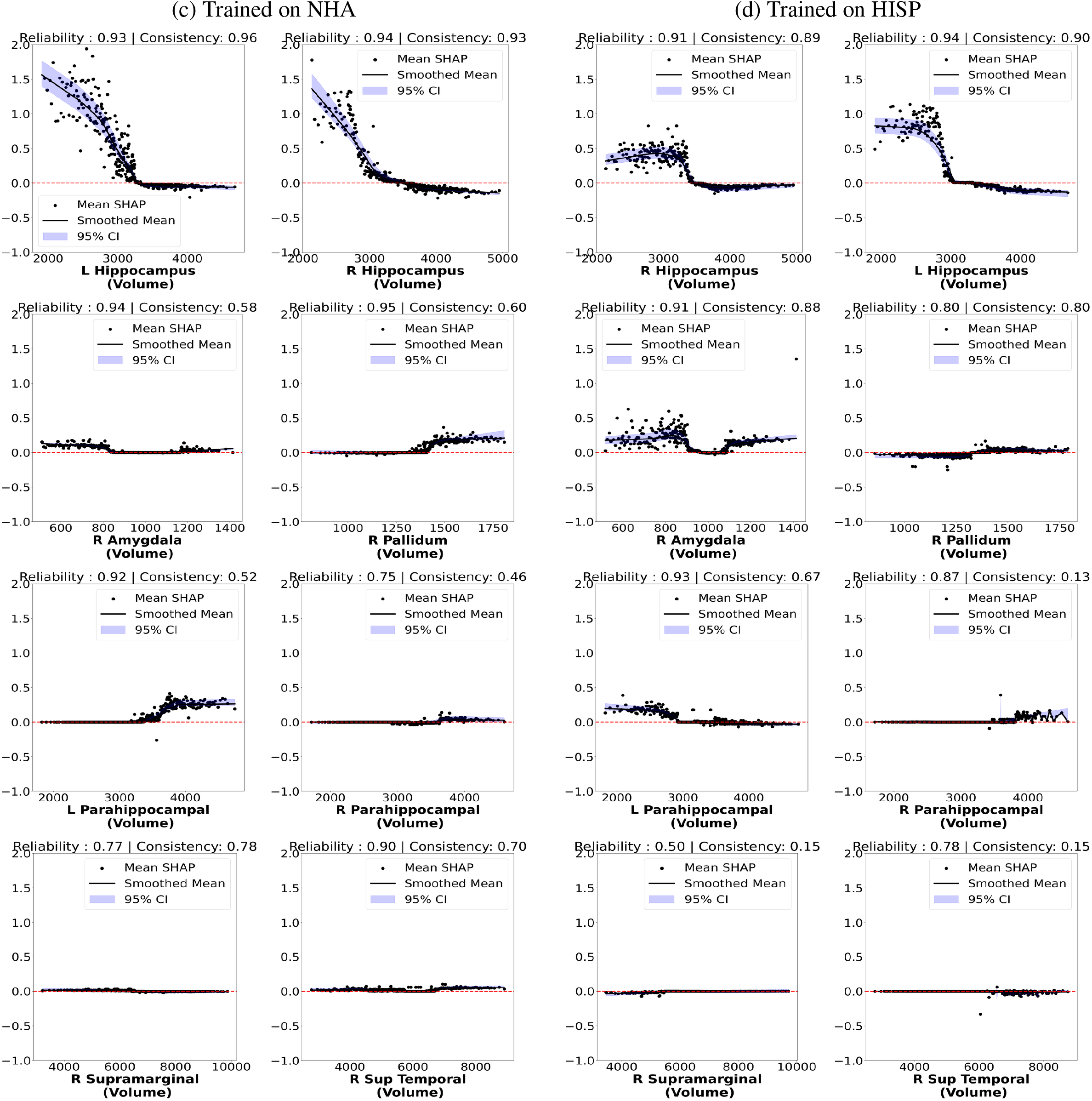
Partial dependence plots of eight dementia-related brain regions guided by ALE meta-analysis across models trained on different populations (ALL and NHW). Each plot displays regional brain volume (x-axis) against SHAP values (y-axis). Where ALE : activation likelihood estimation · SHAP : SHapley Additive exPlanations · CI : confidence interval · ALL : all racial/ethnic populations · NHW : Non-Hispanic White · L : left · R : right · Sup: suprior. Partial dependence plots of eight dementia-related brain regions guided by ALE meta-analysis across models trained on different populations (NHA and HISP). Where ALE : activation likelihood estimation · SHAP : SHapley Additive exPlanations · CI : confidence interval · NHA : Non-Hispanic African · HISP : Hispanic · L : left · R : right · Sup: suprior.

Among the remaining ALE-guided regions, the right amygdala showed modest predictive value, particularly in the All- and NHW-trained models, where lower volumes were associated with an increased risk of dementia. In contrast, the right pallidum demonstrated slightly weaker SHAP gradients in the four models, indicating less classically associated with dementia classification. Furthermore, the parahippocampal gyrus exhibited weak and variable SHAP contributions across all training scenarios, indicating a limited diagnostic influence compared to the hippocampus. Interestingly, the left parahippocampal gyrus exhibited contrasting SHAP contribution trends between the NHA- and HISP-trained models. In the NHA-trained model, increasing volume in this region was associated with higher SHAP values, indicating greater contributions to dementia prediction with larger volumes. Conversely, the HISP-trained model revealed the opposite pattern, suggesting that reduced volume in the left parahippocampal gyrus was more indicative of dementia in the Hispanic population. In the right supramarginal gyrus and right superior temporal gyrus, the SHAP patterns across most regions were unstable or flat, especially for NHA- and HISP-trained models, indicating limited or ambiguous contributions to dementia prediction and requiring the need for sufficient sample representation to derive meaningful attributions. These findings underscore that SHAP-based partial dependence plots can reveal consistent volume thresholds for dementia risk in well-represented populations, but their diagnostic utility decreases under limited or imbalanced training data.

## Discussion

This study examines the fairness and interpretability of machine learning models for dementia classification using structural MRI data from racially and ethnically diverse populations. Using 3, 127 participants from the NACC cohort, we analyzed diagnostic discrepancies across White American, African American, and Hispanic cohorts and demonstrated that conventional ML models trained on a single population suffer from substantial performance decreases and increased false positive and negative rates when applied to underrepresented groups. To address these discrepancies, we evaluated five bias mitigation strategies, including data harmonization, correlation remover, kernel mean matching, semi-supervised domain adaptation, and a novel method we developed, RegAlign. We further integrated interpretable AI techniques using SHAP to identify population-specific contributions from brain regions and validated these findings against a meta-analysis of VBM studies.

A significant contribution of this work is the development of RegAlign, a regularized few-shot learning framework designed to improve both fairness and predictive accuracy when training data from minoritized groups are sparse. RegAlign incorporates three core components: (1) a focal-weighted loss that prioritizes difficult-to-classify cases in imbalanced datasets, (2) class-sensitive weighting that emphasizes dementia detection in underrepresented samples, and (3) class-conditional alignment to harmonize prediction distributions between source and target domains. By integrating these mechanisms, RegAlign achieved a balanced accuracy of 87.9% and reduced inter-group variation in FNR and FPR to below 2%, outperforming other methods across most configurations (Figures 2 and 3). When trained on individual populations, RegAlign also maintained strong performance, particularly in reducing FPR, making it a robust choice under limited data conditions. While SSDA achieved competitive fairness in some cases, it often overfits the positive class, which could compromise generalizability, especially in underrepresented groups.

SHAP-based interpretation provided further insights into model behavior across different training scenarios, and enhanced the explainability of our findings. As shown in Figures 4 and 5, SHAP-based partial dependence plots revealed biologically plausible attribution patterns, particularly for the bilateral hippocampus regions, which are known to experience early atrophy in Alzheimer’s disease. Models trained on all populations and the NHA group consistently displayed steep SHAP transitions between 3000–3500 mm^3^ in hippocampal volume, suggesting data-driven thresholds for dementia risk stratification. In contrast, the attribution patterns in the HISP-trained model were more variable or flat across most brain regions, reflecting the challenges of training on underrepresented subgroups. Among other ALE-guided regions, such as the right amygdala, predictive contributions were modest and more sensitive to population-specific training. These findings suggest that SHAP-based dependence curves may be helpful in deriving interpretable volume thresholds, however, their diagnostic utility could be affected by imbalanced or sparse data.

In general, these findings suggest that AI models require both methodological discrepancy mitigation methods and inclusive training datasets to achieve fairness. While discrepancy mitigation methods, such as RegAlign, can substantially reduce performance gaps, training on representative and demographically diverse cohorts remains essential for reliable and fair dementia diagnosis. Future work should extend these findings to other neurodegenerative conditions, integrate longitudinal trajectories, and explore fairness across sex, age, and socioeconomic factors to further advance responsible and representative AI in healthcare.

## Data and Code Availability

The data used in this study was provided by the NACC (National Alzheimer’s Coordinating Center) cohort (http://www.alz.washingfaton.edu/). The list of VBM studies combined to produce the meta-analysis map is provided in Supplementary materials (Table S2). All source data are available online in the BrainMap VBM Database (Vanasse et al., 2018) at BrainMap.org and portal.BrainMap.org. The code supporting this study is available at: https://github.com/UTHSCSA-NAL/explainable_and_fair_ML_for_ADRD

## Acknowledgements

This study was supported in part by the National Institute of Health (NIH) grant P30AG066546 (South Texas Alzheimer’s Disease Research Center) and grant number 1U24AG074855, 5R01AG080 1R01AG085571, 5R01AG083865, and MH074457. The NACC database is funded by NIA/NIH Grant U24 AG072122. NACC data are contributed by the NIA-funded ADRCs: P30 AG062429 (PI James Brewer, MD, PhD), P30 AG066468 (PI Oscar Lopez, MD), P30 AG062421 (PI Bradley Hyman, MD, PhD), P30 AG066509 (PI Thomas Grabowski, MD), P30 AG066514 (PI Mary Sano, PhD), P30 AG066530 (PI Helena Chui, MD), P30 AG066507 (PI Marilyn Albert, PhD), P30 AG066444 (PI John Morris, MD), P30 AG066518 (PI Jeffrey Kaye, MD), P30 AG066512 (PI Thomas Wisniewski, MD), P30 AG066462 (PI Scott Small, MD), P30 AG072979 (PI David Wolk, MD), P30 AG072972 (PI Charles DeCarli, MD), P30 AG072976 (PI Andrew Saykin, PsyD), P30 AG072975 (PI David Bennett, MD), P30 AG072978 (PI Neil Kowall, MD), P30 AG072977 (PI Robert Vassar, PhD), P30 AG066519 (PI Frank LaFerla, PhD), P30 AG062677 (PI Ronald Petersen, MD, PhD), P30 AG079280 (PI Eric Reiman, MD), P30 AG062422 (PI Gil Rabinovici, MD), P30 AG066511 (PI Allan Levey, MD, PhD), P30 AG072946 (PI Linda Van Eldik, PhD), P30 AG062715 (PI Sanjay Asthana, MD, FRCP), P30 AG072973 (PI Russell Swerdlow, MD), P30 AG066506 (PI Todd Golde, MD, PhD), P30 AG066508 (PI Stephen Strittmatter, MD, PhD), P30 AG066515 (PI Victor Henderson, MD, MS), P30 AG072947 (PI Suzanne Craft, PhD), P30 AG072931 (PI Henry Paulson, MD, PhD), P30 AG066546 (PI Sudha Seshadri, MD), P20 AG068024 (PI Erik Roberson, MD, PhD), P20 AG068053 (PI Justin Miller, PhD), P20 AG068077 (PI Gary Rosenberg, MD), P20 AG068082 (PI Angela Jefferson, PhD), P30 AG072958 (PI Heather Whitson, MD), P30 AG072959 (PI James Leverenz, MD).

