## supplementary materials for "Advancing Fair and Explainable Machine Learning for Neuroimaging Dementia Pattern Classification in Multi-Ethnic Populations"

### Representative and Explainable AI for Multi-Ethnic Dementia Pattern Recognition.

#### Experimental settings

Hyperparameter tuning was performed using GridSearchCV with 10-fold cross-validation to optimize the performance of the XGBoost classifier. This approach systematically evaluated all combinations of specified hyperparameter values to identify the configuration that achieved the best classification results. Table S1 details the default settings and the search ranges used for optimization in this study.

**Table S1:** Default and tuning parameters for hyperparameter optimization. Default values serve as the initial settings, while the tuning grid defines the range of values explored during GridSearchCV with 10-fold cross-validation.

| Method | Parameter | Default value | Tuning values | Description |
|---|---|---|---|---|
| XGBoost | Max depth | 2 | $[2, 3, 4, 5, 6]$ | Maximum depth of a tree; controls model complexity and overfitting. |
| | Learning rate | 1 | $[0.1, 0.3, 0.5, 0.7, 1]$ | Step size shrinkage to prevent overfitting and improve convergence. |
| | Lambda | 1 | $[1, 3, 5, 7]$ | L2 regularization term on weights to reduce overfitting. |
| | Alpha | 0 | $[0, 1, 3, 5]$ | L1 regularization term on weights to encourage sparsity. |
| | Scale positive weight | 1 | $[1, 1.15, 1.3, 1.45]$ | Balances the dataset by scaling the weight of positive examples. |
| SVM | C | 1 | $[0.001, 0.01, 0.1, 1, 10]$ | Regularization parameter; controls the trade-off between margin width and classification error. |
| | Kernel | `rbf` | {`rbf`, `linear`} | Specifies the kernel type used in the algorithm. |

The training process followed two distinct strategies to evaluate model performance and fairness across populations, as illustrated in Figure 1 (Panel C), Main Manuscript. In the first strategy, the entire dataset, comprising samples from all racial and ethnic groups, was partitioned into 10 folds for outer cross-validation to estimate general performance. For each outer fold, we performed a 10-fold inner cross-validation defined by GridSearchCV on the 9 training folds to identify the optimal hyperparameters ($1,600$ combinations for XGBoost and $10$ for SVM). After selecting the best-performing configuration, the model was retrained on all $9$ folds and evaluated on the held-out test fold. This process was repeated across all $10$ folds, ensuring robust performance estimation and exposing the model to diverse population characteristics during training and improve model generalization.

In the second strategy, the dataset was filtered to include only a specific population group for training, and similarly partitioned into $10$ folds. For each outer fold, the model processed the same grid search and cross-validation process as above. However, evaluation was conducted not only on the held-out fold from the same population but also on data from all other populations that were not seen during training. This approach enabled us to quantify how well models trained on a specific group generalized to out-of distribution populations. Both strategies provided a comprehensive assessment of model performance and fairness across demographic groups.

#### Reliability and Consistency Metrics

To assess the robustness of SHAP value estimates across $5,000$ bootstrap models, we quantified two key metrics: reliability and consistency.

##### Reliability

The reliability metric captures the dispersion of average absolute SHAP values across bootstrap replicates using the coefficient of variation. For each bootstrap replicate $i$, let $\mu_i$ be the mean absolute SHAP value. The reliability is computed as:

$$\text{Reliability} = \max\left(0,\ 1 - \min\left(1,\ \frac{\sigma_\mu}{\bar{\mu} + \epsilon}\right)\right), \tag{S1}$$

where $\bar{\mu} = \frac{1}{N}\sum_{i=1}^{N} \mu_i$ is the mean of the SHAP magnitudes, $\sigma_\mu$ is their sample standard deviation, $N$ is the number of bootstrap replicates, and $\epsilon$ is a small constant (e.g., $10^{-8}$) to avoid division by zero.

##### Consistency

To evaluate directional agreement across bootstrap replicates, we define the consistency score. Let $v_{ij}$ denote the SHAP value in volume bin $j$ from replicate $i$, and $s_{ij} = \text{sign}(v_{ij})$. Then the average sign across $N$ replicates is:

$$\bar{s}_j = \frac{1}{N}\sum_{i=1}^{N} s_{ij}, \tag{S2}$$

and the consistency is defined as the fraction of bins where the sign is strongly aligned:

$$\text{Consistency} = \frac{1}{B}\sum_{j=1}^{B} \mathbb{I}\left(|\bar{s}_j| > \theta_c\right), \tag{S3}$$

where $B$ is the number of bins, $\mathbb{I}(\cdot)$ is the indicator function that equals 1 when the condition is satisfied and 0 otherwise, and $\theta_c$ is the directional agreement threshold. In our experiments, we set $\theta_c = 0.7$, which serves as a practical cutoff indicating that at least 70% of bootstrap replicates agree on the direction of SHAP values in a given bin.

#### VBM studies included in the meta-analysis

In particular, the preliminary meta-analysis map was generated by reprocessing a set of voxel-based morphometry (VBM) studies collected from a previous meta-analysis [36]. We created a brain map by applying the activation likelihood estimation (ALE) method, using the GingerALE software (version 3.0.2) [11], to combine the selected VBM studies into a single ALE map that highlights brain regions where atrophy is likely associated with Alzheimer's disease versus Normal Cognition. Following the procedures outlined in [36], the ALE analysis was conducted using a cluster-forming threshold of $0.001$ and a cluster-level significance threshold of $0.05$, with statistical significance estimated through $1,000$ random permutations. The continuous ALE map produced by GingerALE, in which non-significant brain areas were assigned null values, was then thresholded at the smallest non-zero value to create a binary map, facilitating comparisons across different training strategies. Finally, the ALE meta-analysis identified several brain regions with statistically significant activation likelihood associated with dementia. The $38$ voxel-based morphometry (VBM) studies with $47$ experiments combined in our meta-analysis are reported in Table S2.

#### Performance on Dementia Classification using SVM

We conducted additional experiments using alternative classifiers, such as Support Vector Machines (SVM), to further examine classification discrepancies. Since SVMs do not support custom objective functions, we applied only data harmonization, correlation regularization (CR), and kernel mean matching (KMM) for discrepancy mitigation. As shown in Figure S1, training on data from all populations yielded the highest overall accuracy, reaching up to $83.5\%$, and consistently reduced performance gaps across populations. Regarding FPR gaps when trained on individual populations, data harmonization produced the smallest gap for NHW and NHA, while KMM was most effective for the Hispanic group. In terms of FNR gaps, KMM performed best when trained on NHW, CR yielded the smallest gap for NHA, and the baseline SVM model achieved the lowest FNR gap when trained on Hispanic data. These results indicate that diagnosis discrepancies exist when using SVMs, but combining SVMs with appropriate discrepancy mitigation techniques, particularly tailored to each population, can slightly reduce group-level performance gaps, although not as effectively as models that support custom objectives.

In addition, Figure S2 illustrates the Pareto front optimization to evaluate the trade-off between classification performance and fairness using SVM classifier. In the top-left panel (BA vs. FPR across methods), data harmonization achieved slightly better trade-offs than other approaches, maintaining low FPR while preserving accuracy. The top-right panel (BA vs. FPR across training scenarios) showed that models trained on NHW achieved the most favorable balance, closely followed by those trained on all populations, highlighting the stability of these training settings. In the bottom-left panel (BA

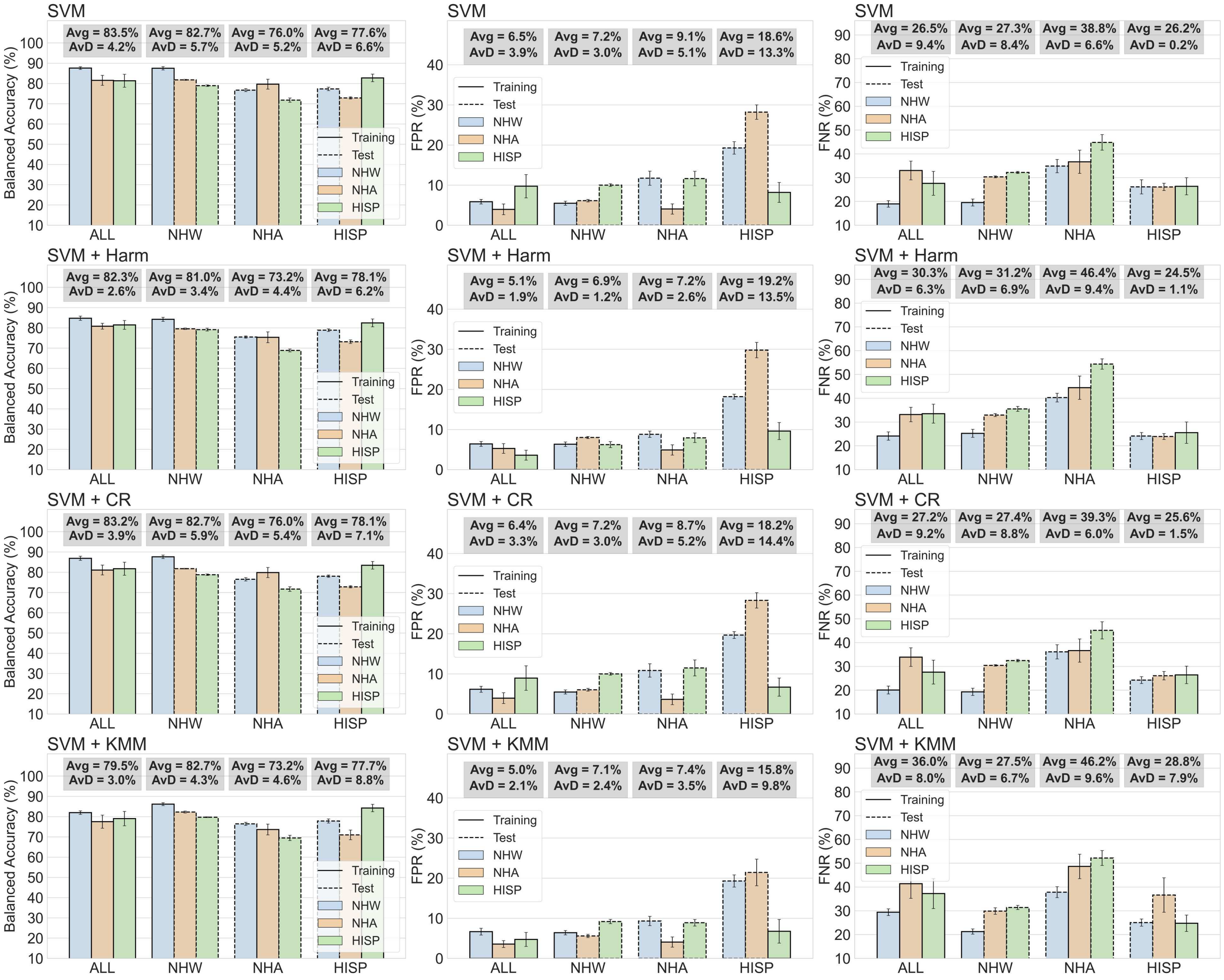


**Figure S1:** Comparison of baseline classifier and three discrepancy mitigation methods across three evaluation metrics: Balanced Accuracy, FPR, and FNR. The four scenarios include training on all racial/ethnic groups (ALL), or training exclusively on NHW, NHA, and HISP group. Where SVM : Support Vector Machine · Harm: data harmonization · CR : correlation remover · KMM kernel mean matching

vs. FNR across methods), both the baseline and CR-based models demonstrated the best trade-offs, minimizing false negatives while sustaining competitive accuracy. Finally, the bottom-right panel (BA vs. FNR across training scenarios) indicated that the model trained on the HISP population achieved the optimal trade-off, though with greater variability, while NHW- and ALL-trained models offered more stable performance with slightly lower fairness. These results emphasize that both training diversity and appropriate mitigation strategies influence the accuracy–fairness balance.

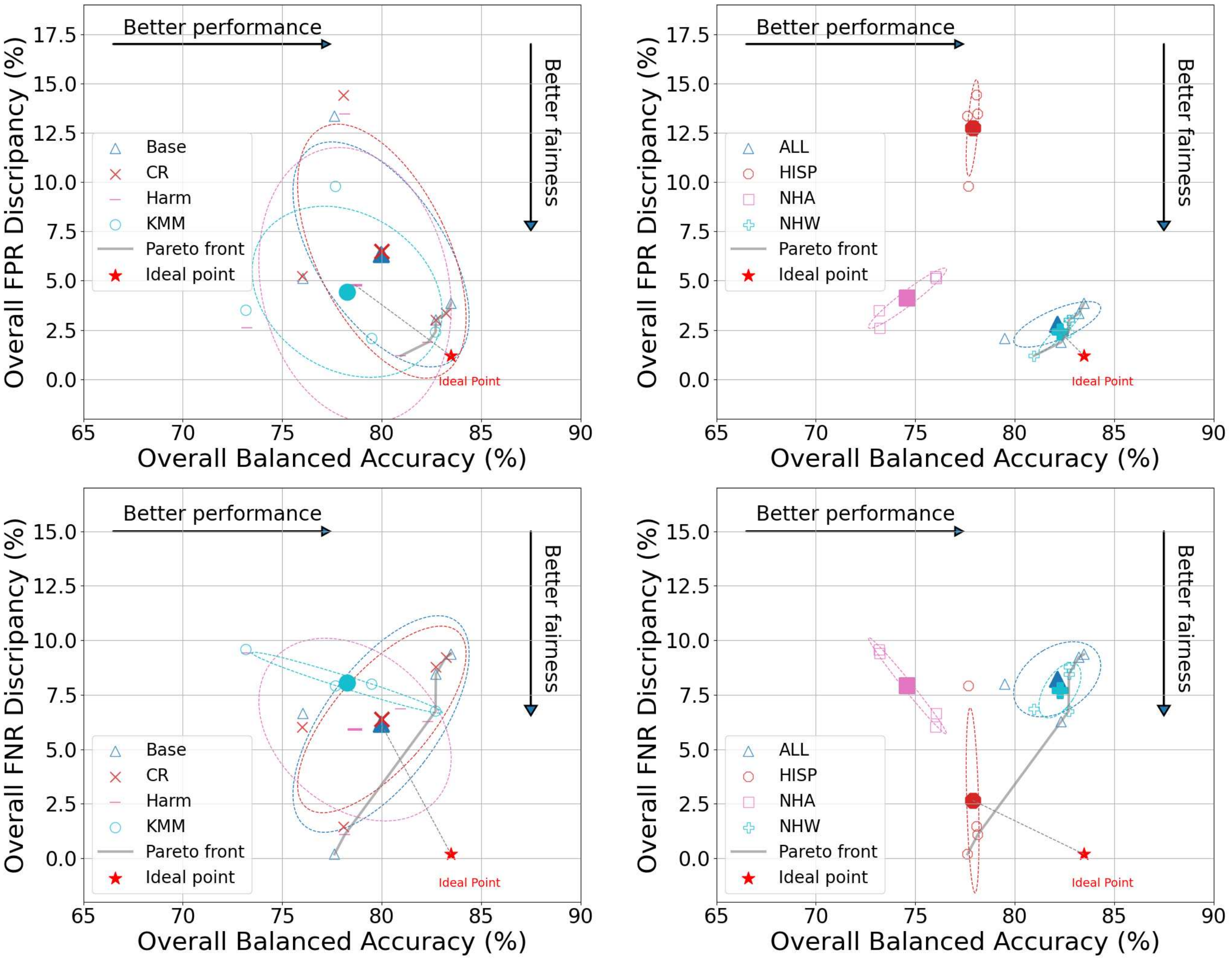


**Figure S2:** Pareto front optimization for balancing performance and fairness based on FPR / FNR discrepancy. Ideal point is defined by the interaction of the highest performance and the lowest fairness. The bigger solid marker denotes the centroid of each group of method or training scenario, the gray dashed line indicates the shortest distance from a centroid to the ideal point, and the ellipse represents the group confidence distribution. Where Harm: data harmonization · CR : correlation removal · KMM kernal mean matching ·.

### Significance Assessment via Bootstrapped Confidence Intervals

To evaluate the reliability of individual brain region contributions, we estimated $95\%$ confidence intervals (CIs) of SHAP values using empirical percentiles over $5,000$ bootstrapping iterations. A region was considered statistically significant if its CI did not cross zero, indicating a consistent direction of effect on model prediction. Confidence interval plots were generated separately for each of the four training scenarios: models trained on all population (**ALL**), **NHW**, **NHA**, and **HISP** populations, as shown in Figure S3. For each scenario, SHAP values were computed across $144$ brain regions, grouped into eight anatomical categories: ventricles ($6$ regions), white matter ($19$), frontal lobe ($36$), occipital lobe ($18$), subcortical structures ($16$), temporal lobe ($10$), parietal and cerebellar regions ($8$), and an additional group labeled "Others" ($31$) encompassing remaining regions. Applying the CI-based significance criterion, the number of regions with non-zero-crossing intervals varied by training cohort: $124$ regions for

**(a)** Trained on ALL

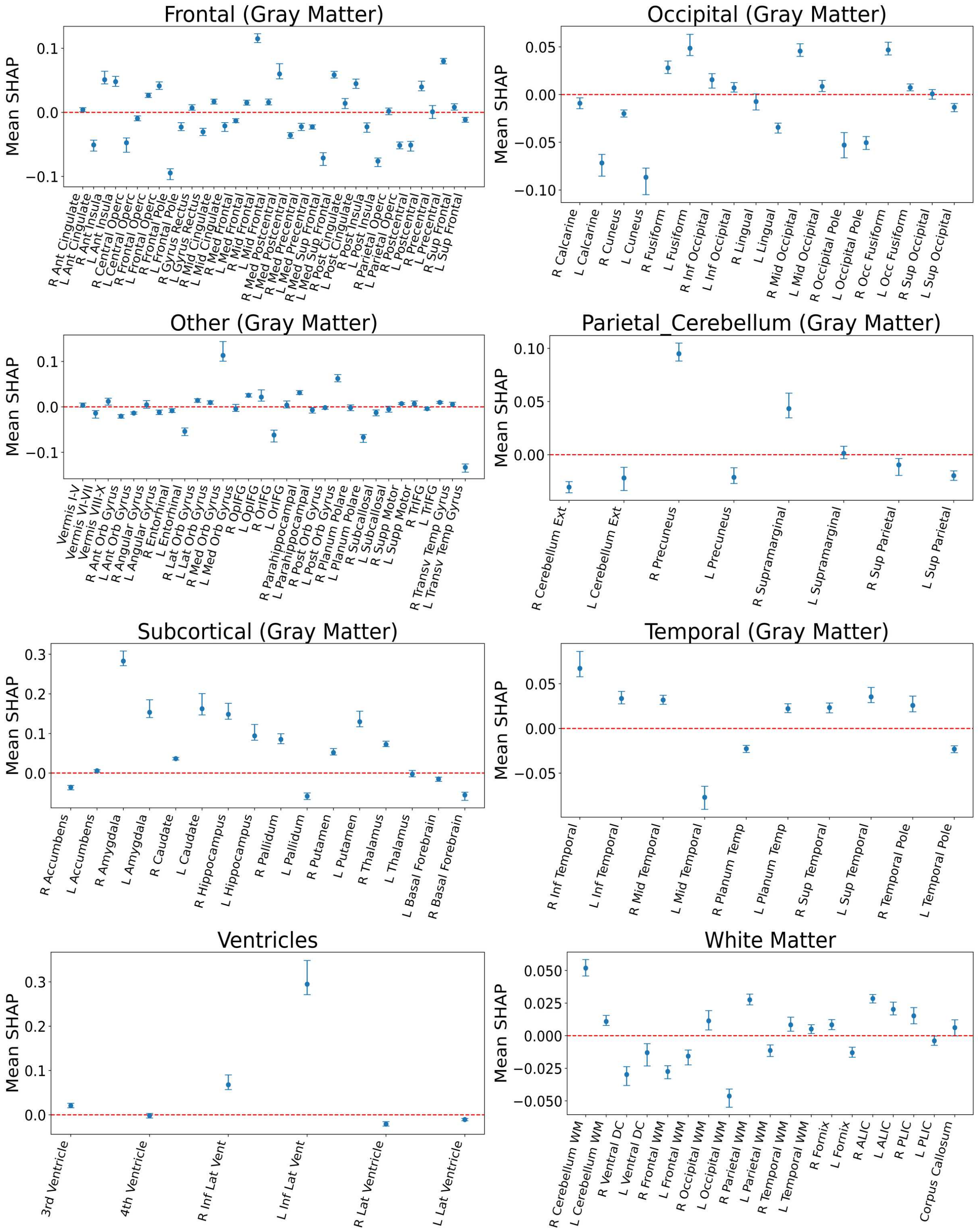


**Figure S3:** Bootstrapped 95% confidence interval plots of SHAP values for the All-trained model across eight brain region groups. Only regions whose CIs did not cross zero-line are selected.

**(b)** Trained on NHW

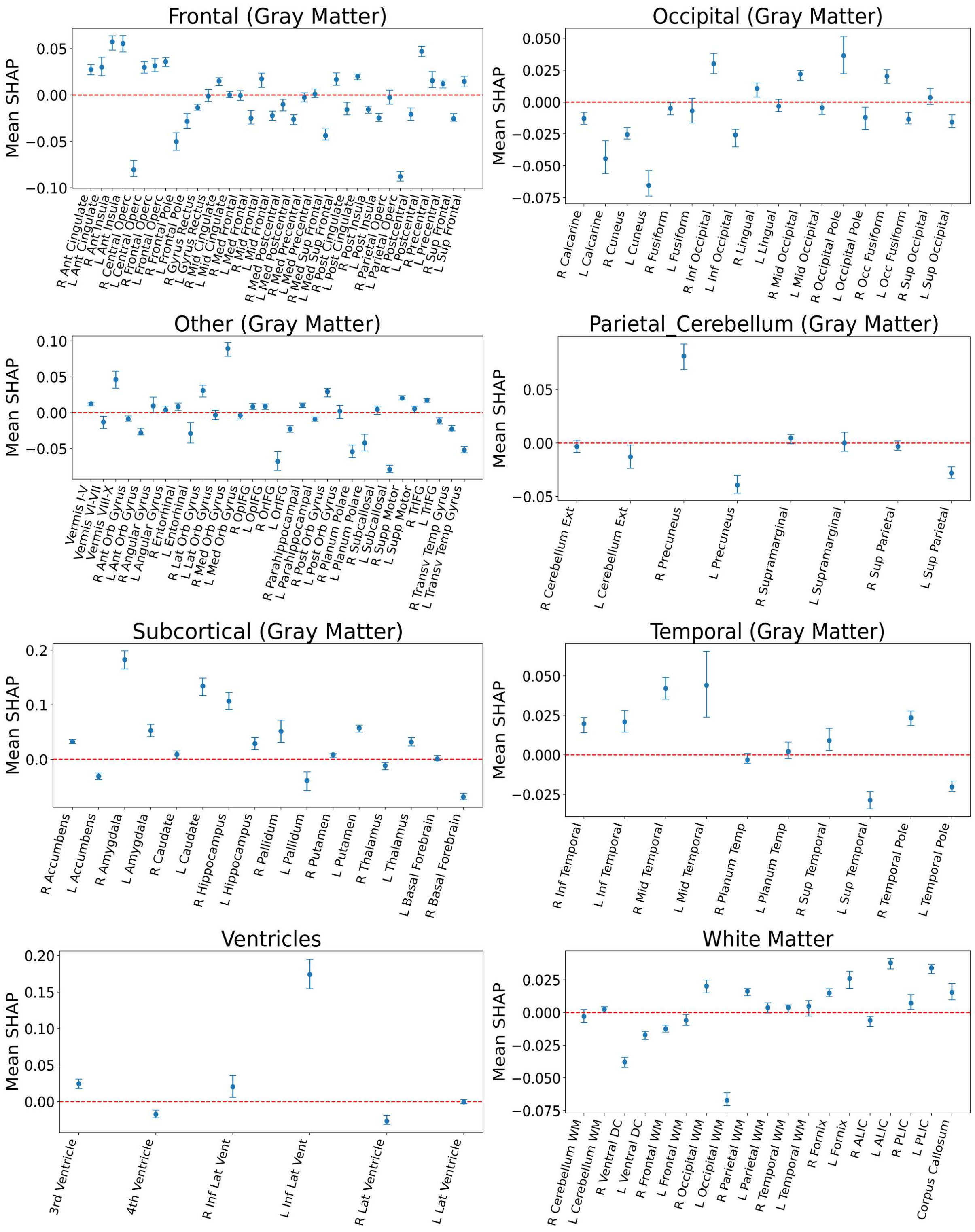


**Figure S3:** (continued) NHW-trained model. Only regions whose CIs did not cross zero-line are selected.

**(c)** Trained on NHA

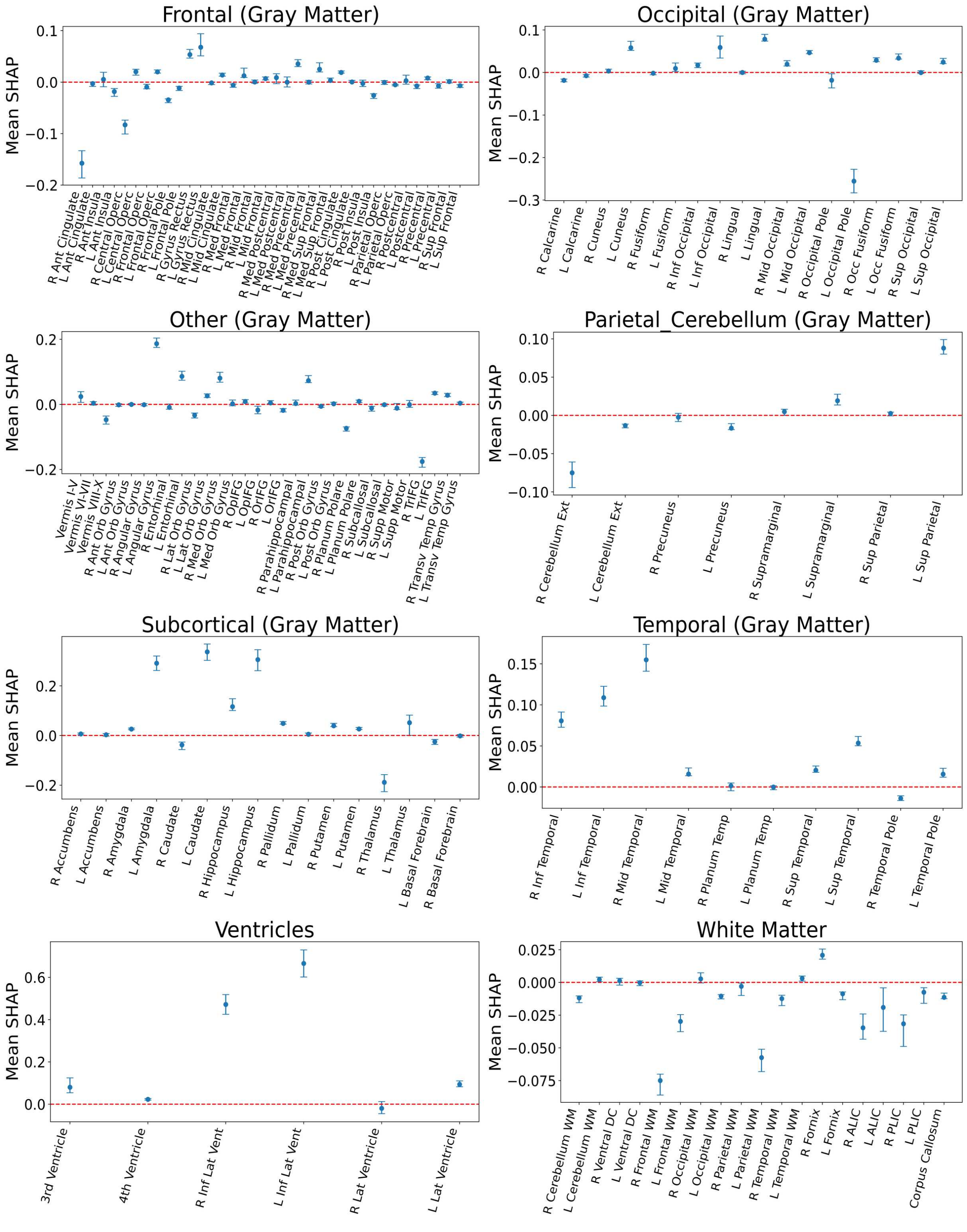


**Figure S3:** (continued) NHA-trained model. Only regions whose CIs did not cross zero-line are selected.

**(d)** Trained on HISP

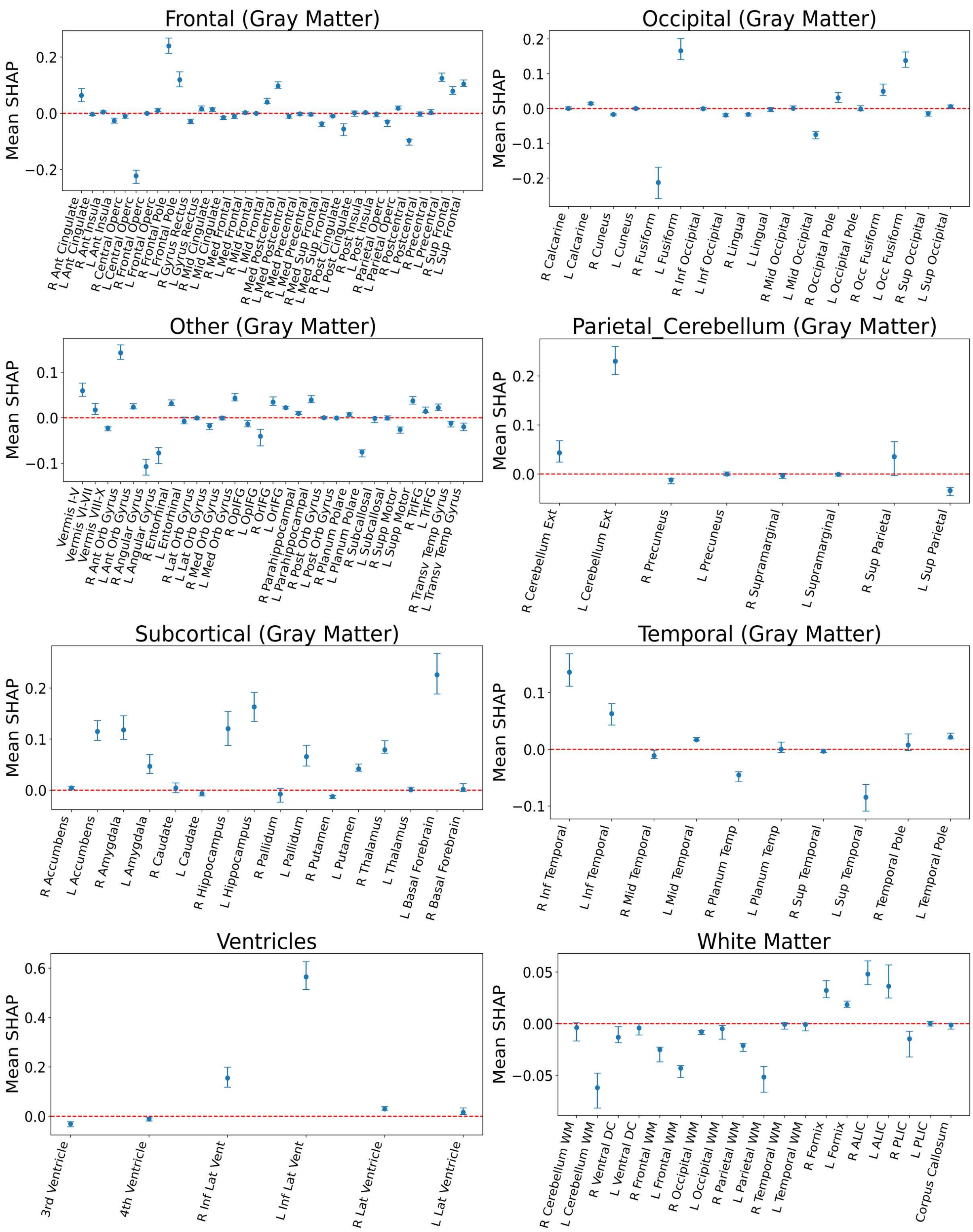


**Figure S3:** (continued) HISP-trained model. Only regions whose CIs did not cross zero-line are selected.

#### SHAP-based Heatmap Visualization for Dementia Prediction

To better understand the regional contributions underlying model predictions, we visualized the remaining SHAP heatmaps across population-specific training scenarios and modeling configurations.

**(1) Baseline XGBoost SHAP Heatmaps without ALE-based Meta Analysis.** We generated SHAP heatmaps for a baseline XGBoost classifier trained independently on four population-specific datasets (All, NHW, NHA, and HISP), without guidance from ALE-based meta-analysis. For each scenario, heatmaps were plotted across three major brain tissue classes: gray matter, white matter, and ventricles, as shown in Figure S4. We generated SHAP heatmaps for a baseline XGBoost classifier trained independently on four population-specific datasets (All, NHW, NHA, and HISP), with and without guidance from ALE-based meta-analysis. For each scenario, heatmaps were plotted across three major brain tissue classes: gray matter, white matter, and ventricles, as shown in Figure S4 (a), (b), and (c). Figure S4 (d) illustrates the average SHAP maps under ALE meta-analysis across four training scenarios. Each scenario includes paired heatmaps highlighting contributions of brain regions toward predicting dementia. The ALE meta-analysis map is displayed in the upper row as a reference, highlighting consistently reported atrophic regions across VBM studies.

**(2) SHAP Heatmaps with Discrepancy Mitigation Techniques.** To assess how discrepancy mitigation techniques influence regional attributions, we visualized SHAP heatmaps (Figure S5) from models trained with five advanced methods: data harmonization, correlation removal (CR), kernel mean matching (KMM), semi-supervised domain adaptation (SSDA), and the proposed RegAlign. For each method, heatmaps were computed separately for dementia and NC predictions across all four training scenarios, resulting in eight plots per method. These comparisons enable evaluation of attribution stability and discrepancy reduction across learning paradigms.

#### SHAP-Based Partial Dependence Analysis of 144 Brain Regions

This section presents the full set of partial dependence plots for 144 brain regions, illustrating the relationship between regional brain volume (x-axis) and SHAP contribution (y-axis) across the classification model. Positive SHAP values indicate contributions toward dementia prediction, while negative values support normal cognition classification. These plots provide insight into how variations in brain structure influence model predictions and help identify region-specific volume thresholds associated with increased dementia risk. All plots were generated from the baseline model trained on the four scenarios, as shown in Figure S6. The list of regions ordered from left to right and top to bottom is illustrated in Figure S6 (e).

**(a)** Gray Matter

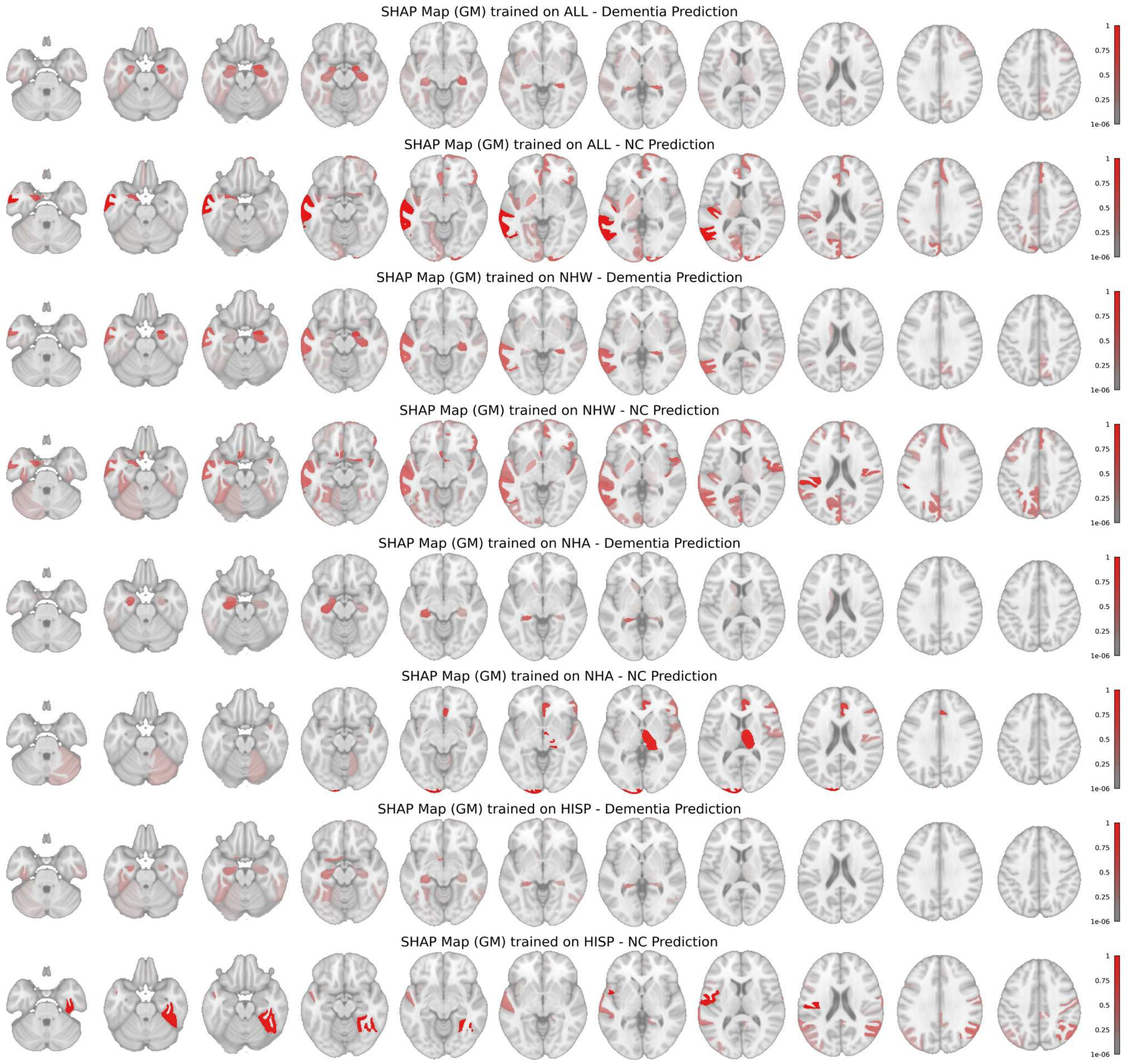


**Figure S4:** SHAP heatmaps without ALE-based meta-analysis of gray matter regions for dementia and NC predictions across four training scenarios using baseline XGBoost. The SHAP maps derived from the baseline model using gray matter features revealed consistent attribution to key dementia-related regions such as the hippocampus during dementia prediction. However, for NC prediction, the patterns appeared scattered and lacked biological coherence, suggesting possible instability in decision-making for non-dementia cases.

**(b)** White Matter

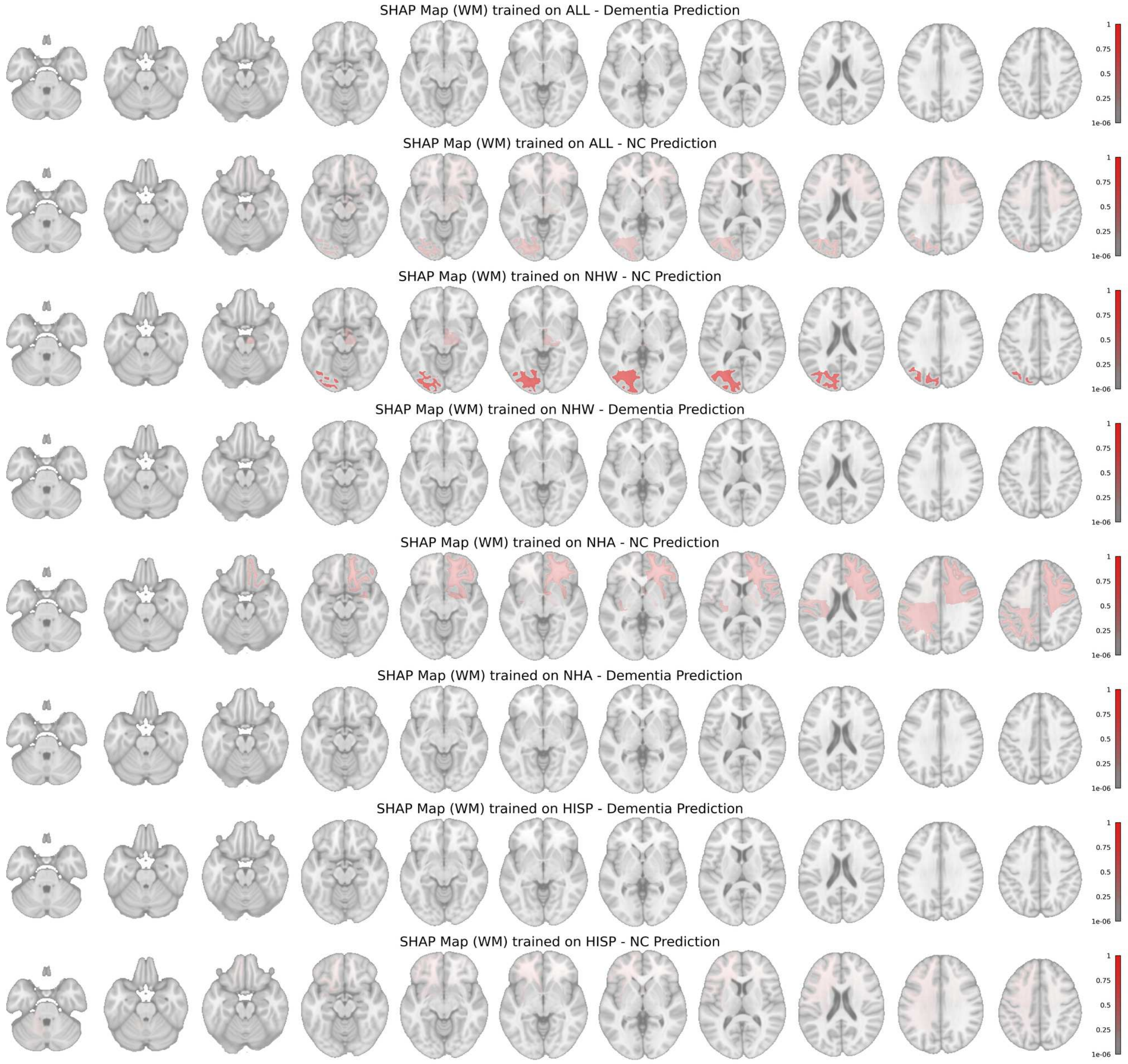


**Figure S4:** (continued) SHAP heatmaps without ALE-based meta-analysis of white matter regions. The SHAP contributions for NC were slightly more structured but overall showed low intensity, indicating limited influence on classification.

**(c)** White Matter

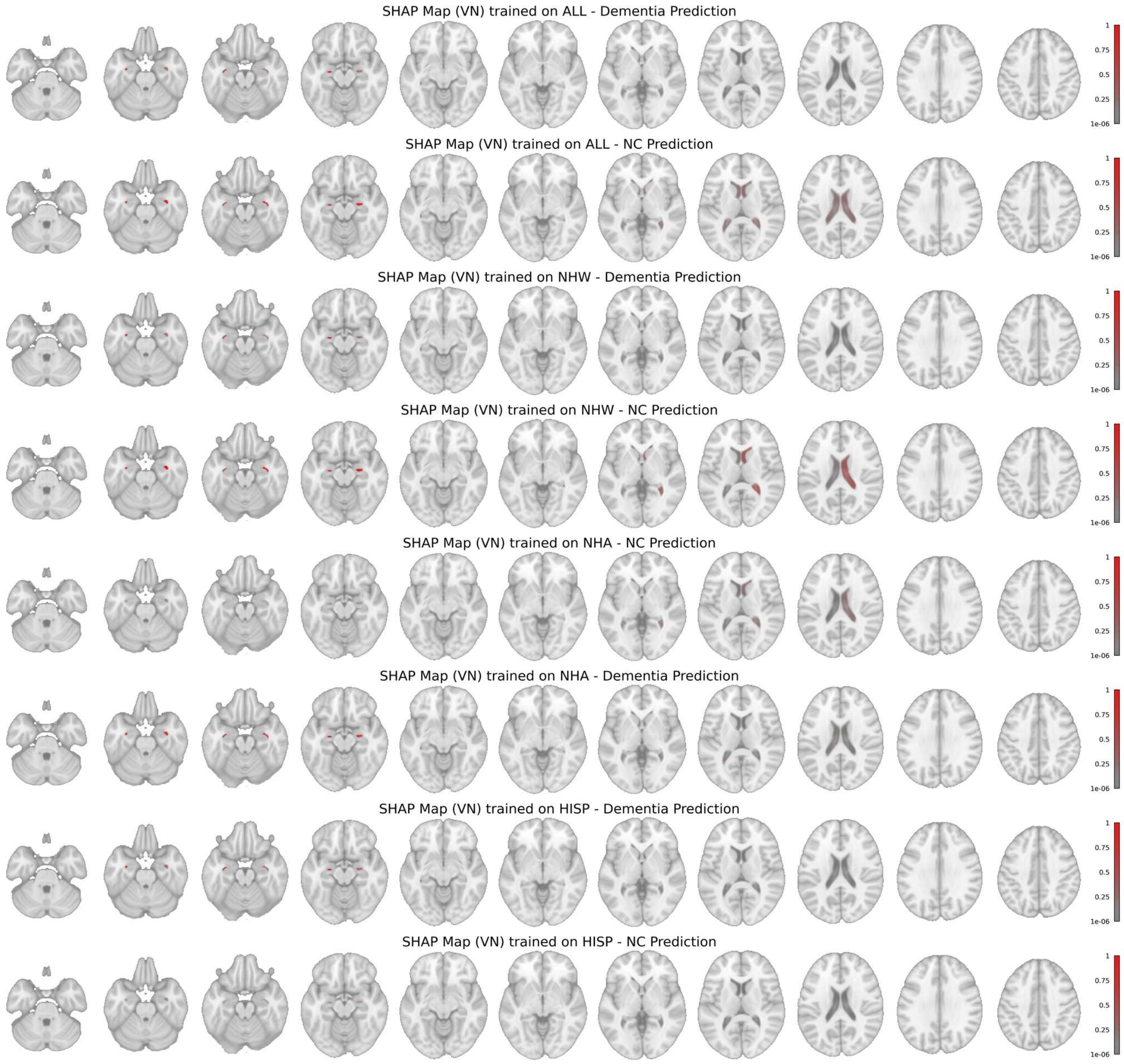


**Figure S4:** (continued) SHAP heatmaps without ALE-based meta-analysis of ventricular regions. The SHAP maps displayed symmetric and interpretable patterns for both dementia and NC predictions when trained on all populations or NHW alone. In contrast, models trained on NHA and HISP produced divergent ventricular attributions across classes, potentially contributing to the higher diagnostic discrepancy observed in these subgroups.

**(d)** ALE-based SHAP Heatmaps

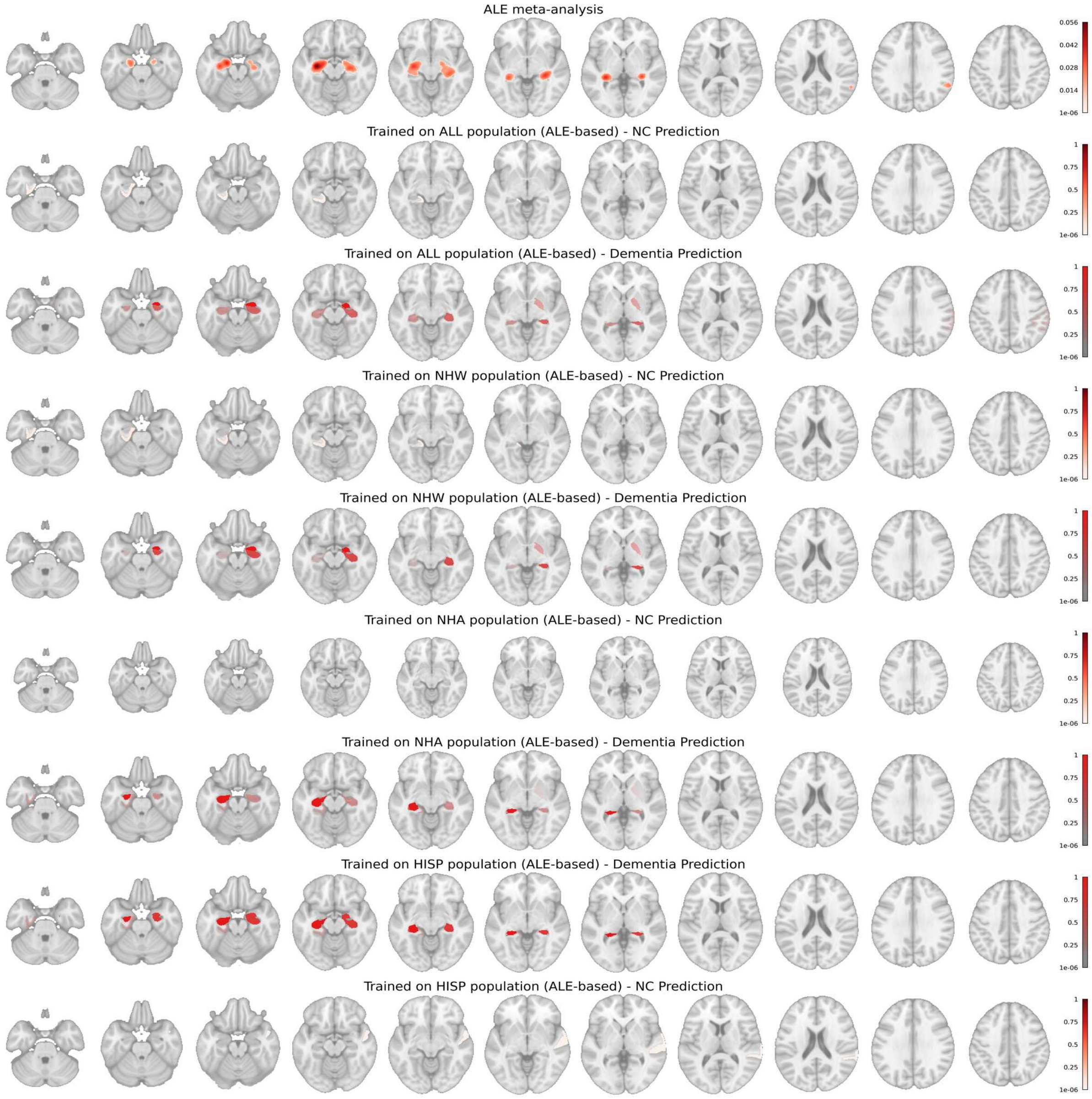


**Figure S4:** (continued) SHAP heatmaps with ALE-based meta-analysis for the four training scenarios. When trained on all or NHW populations, the model predominantly focused on the right hippocampus and parahippocampal gyrus for dementia classification. In contrast, the model trained on NHA data emphasized the left hippocampus, while the HISP-trained model showed bilateral hippocampal regions.

**(a)** Trained on All

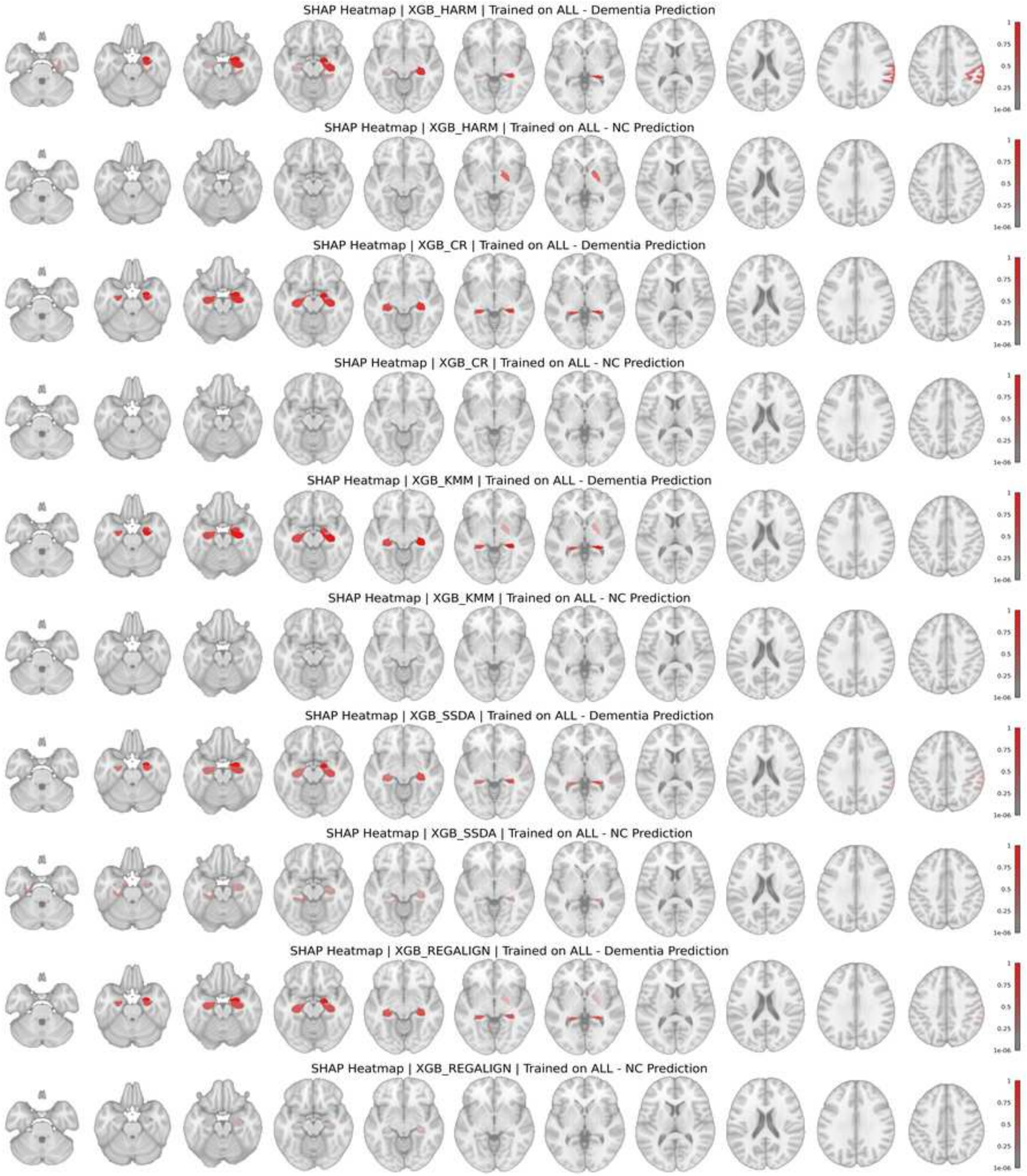


**Figure S5:** SHAP heatmaps with ALE-based meta-analysis of five discrepancy mitigation methods for dementia and NC predictions across all-population training scenario.

**(b)** Trained on NHW

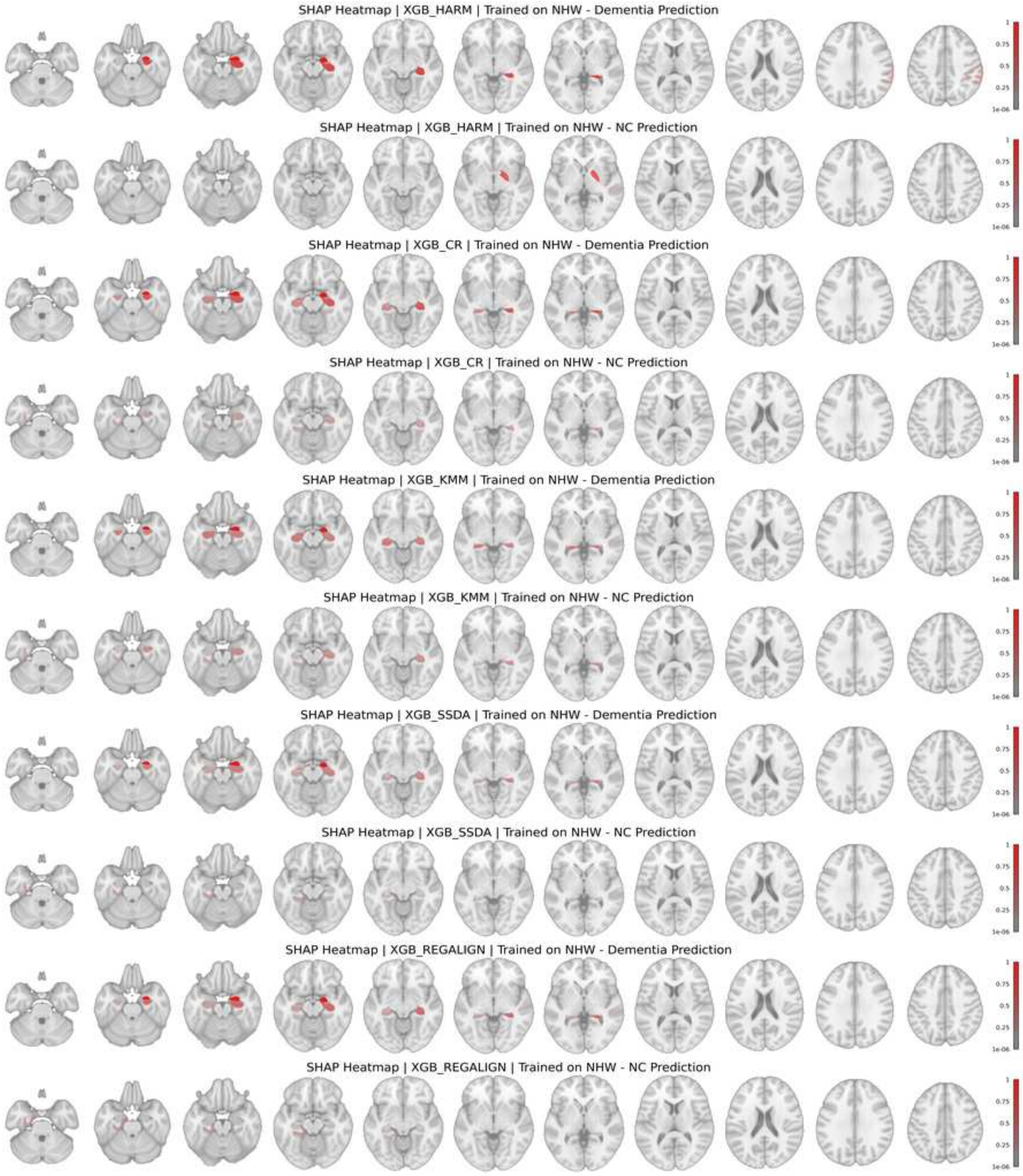


**Figure S5:** (continued) SHAP heatmaps with ALE-based meta-analysis across NHW-population training scenario.

**(c)** Trained on NHA

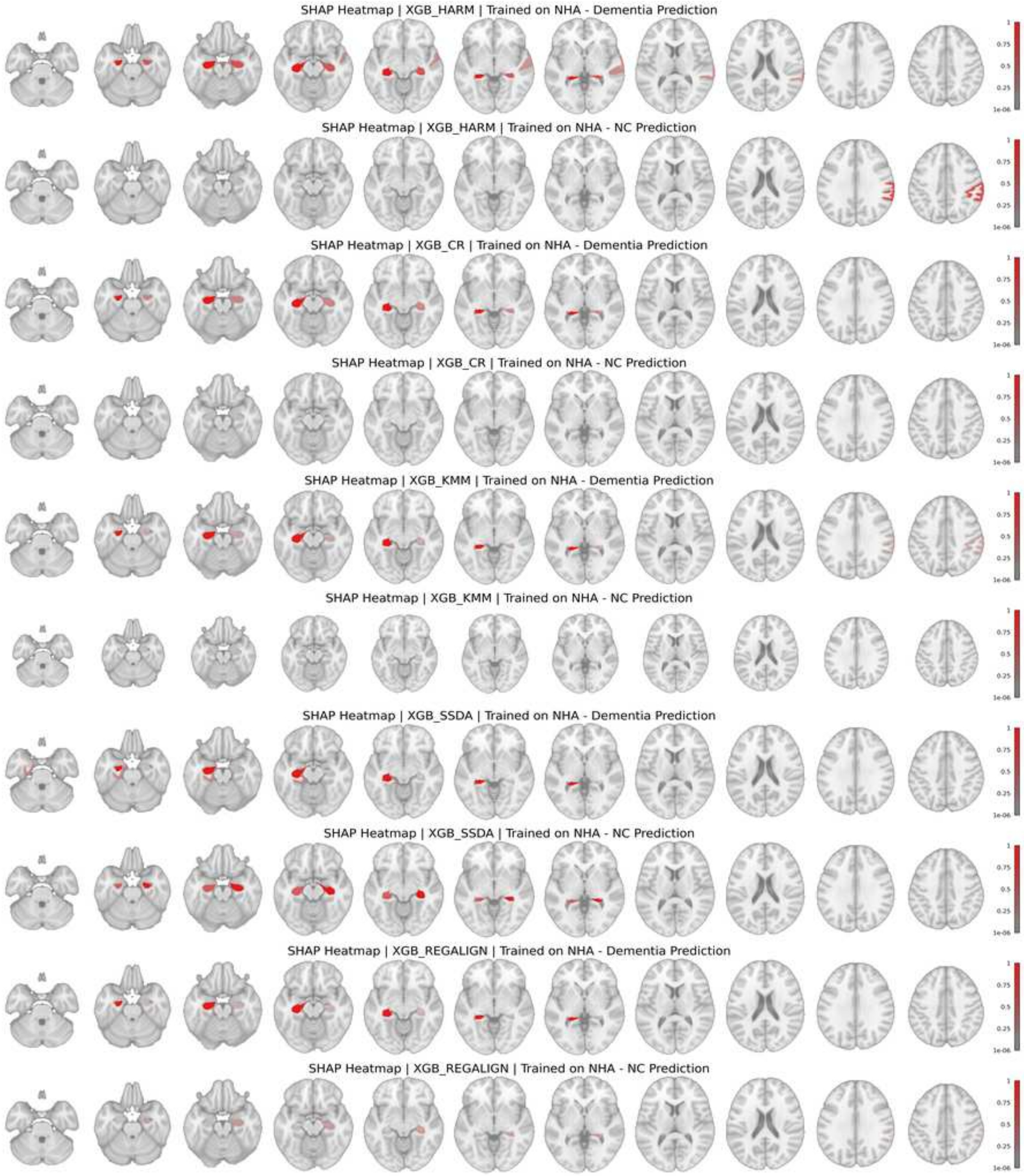


**Figure S5:** (continued) SHAP heatmaps with ALE-based meta-analysis across NHA-population training scenario.

**(d)** Trained on HISP

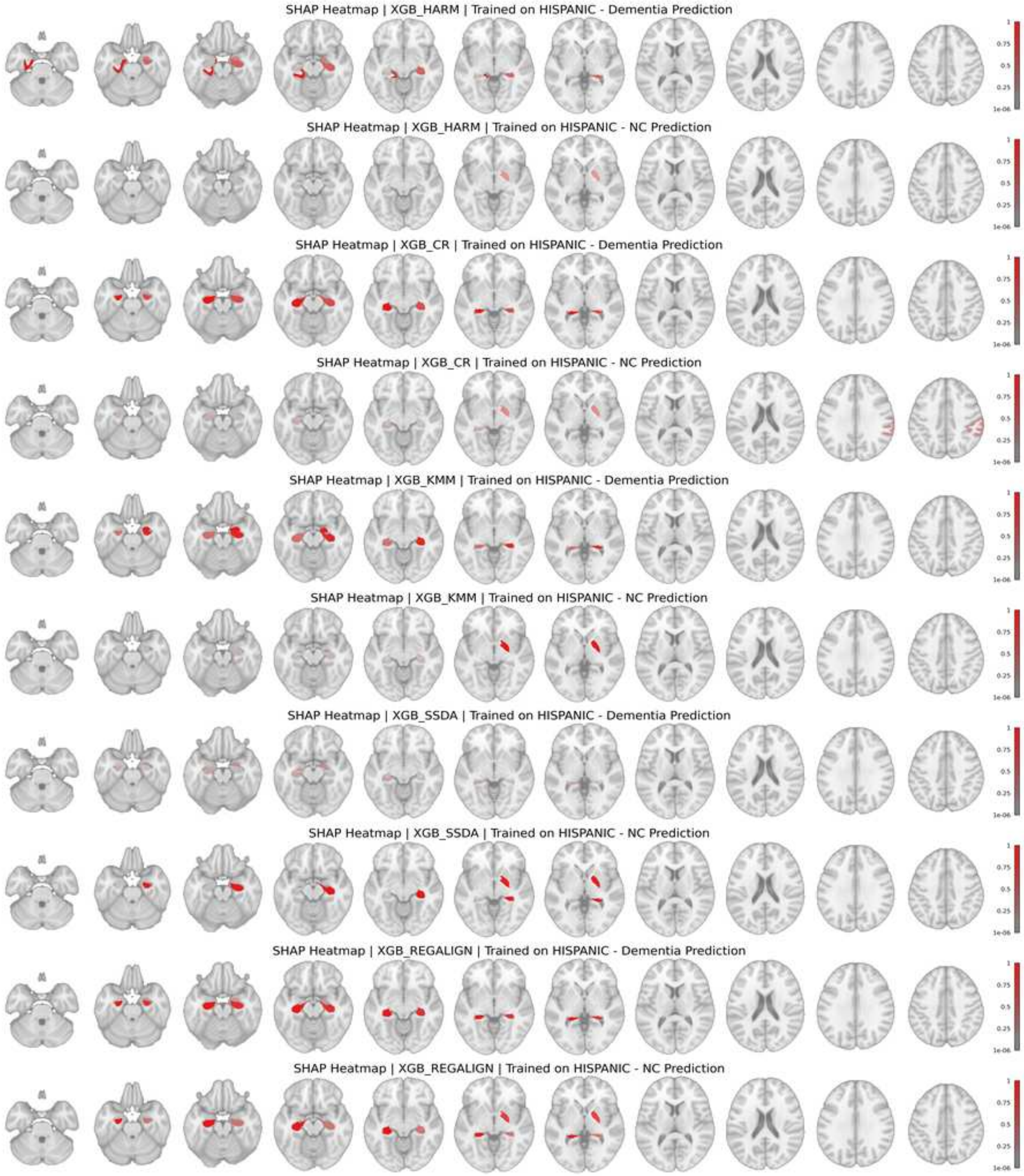


**Figure S5:** (continued) SHAP heatmaps with ALE-based meta-analysis across HISP-population training scenario. SHAP maps generated by the proposed RegAlign revealed consistent attribution patterns for dementia and NC, unlike other methods that exhibited divergent feature usage across classes. Therefore, RegAlign showed superior discrepancy mitigation, achieving both fair and accurate predictions.

**(a)** Trained on ALL

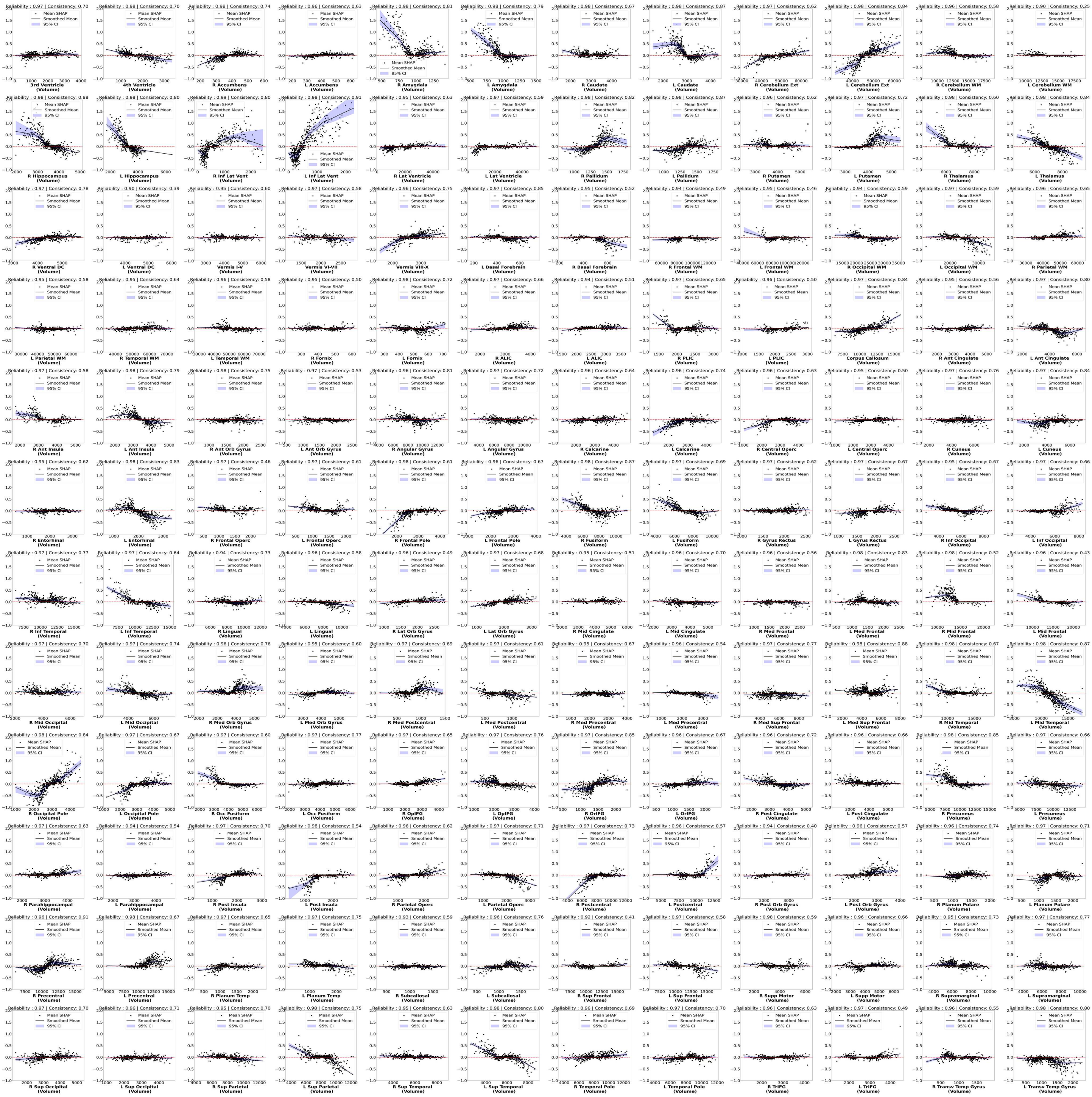

**Figure S6:** Partial dependence plots for all 144 brain regions showing the relationship between regional volume and SHAP contributions for dementia classification. Positive SHAP values indicate contribution toward dementia prediction, while negative values support normal cognition.

(**b**) Trained on NHW

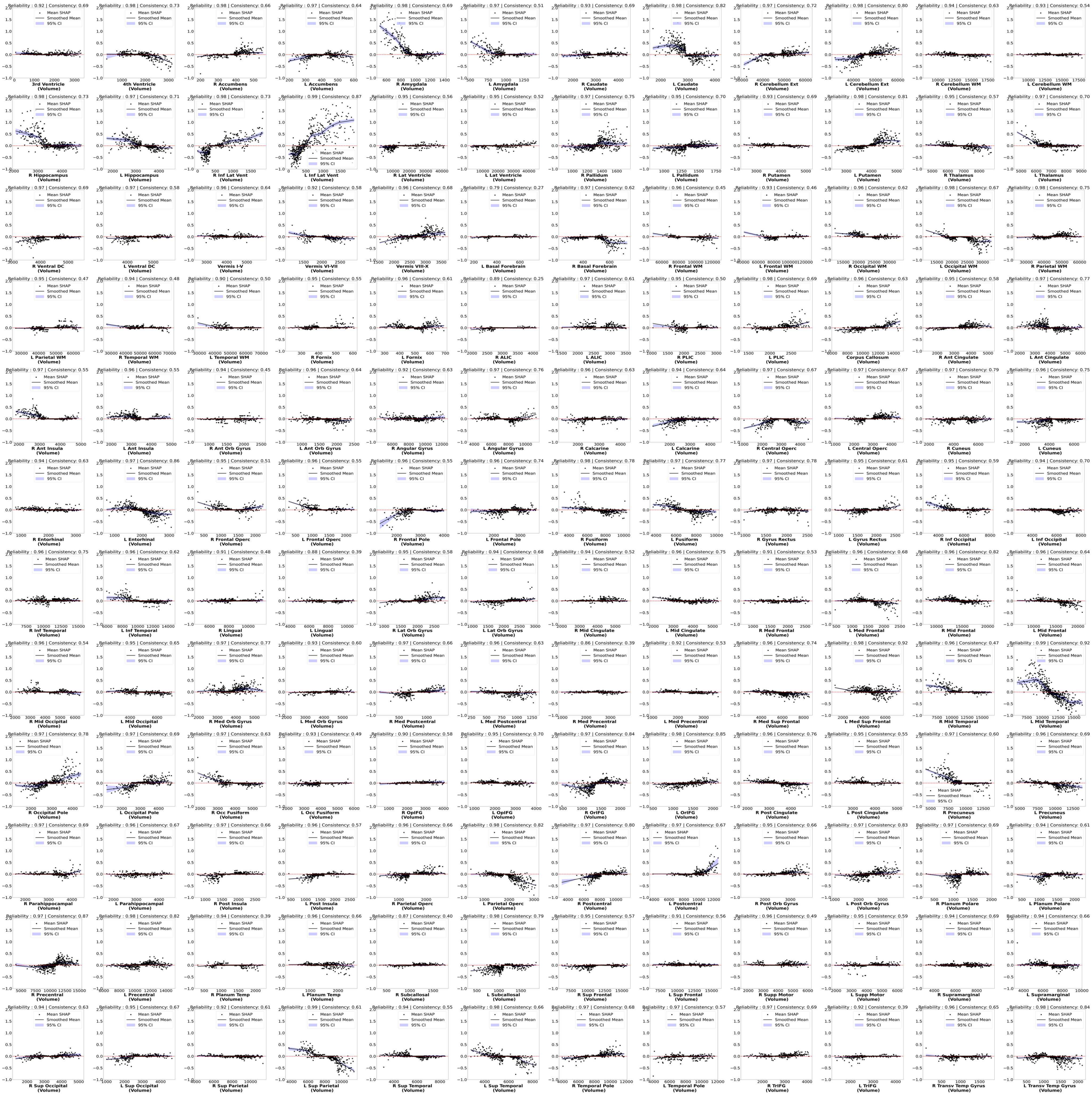

**Figure S6:** (continued) Partial dependence plots for training on NHW.

**(c)** Trained on NHA

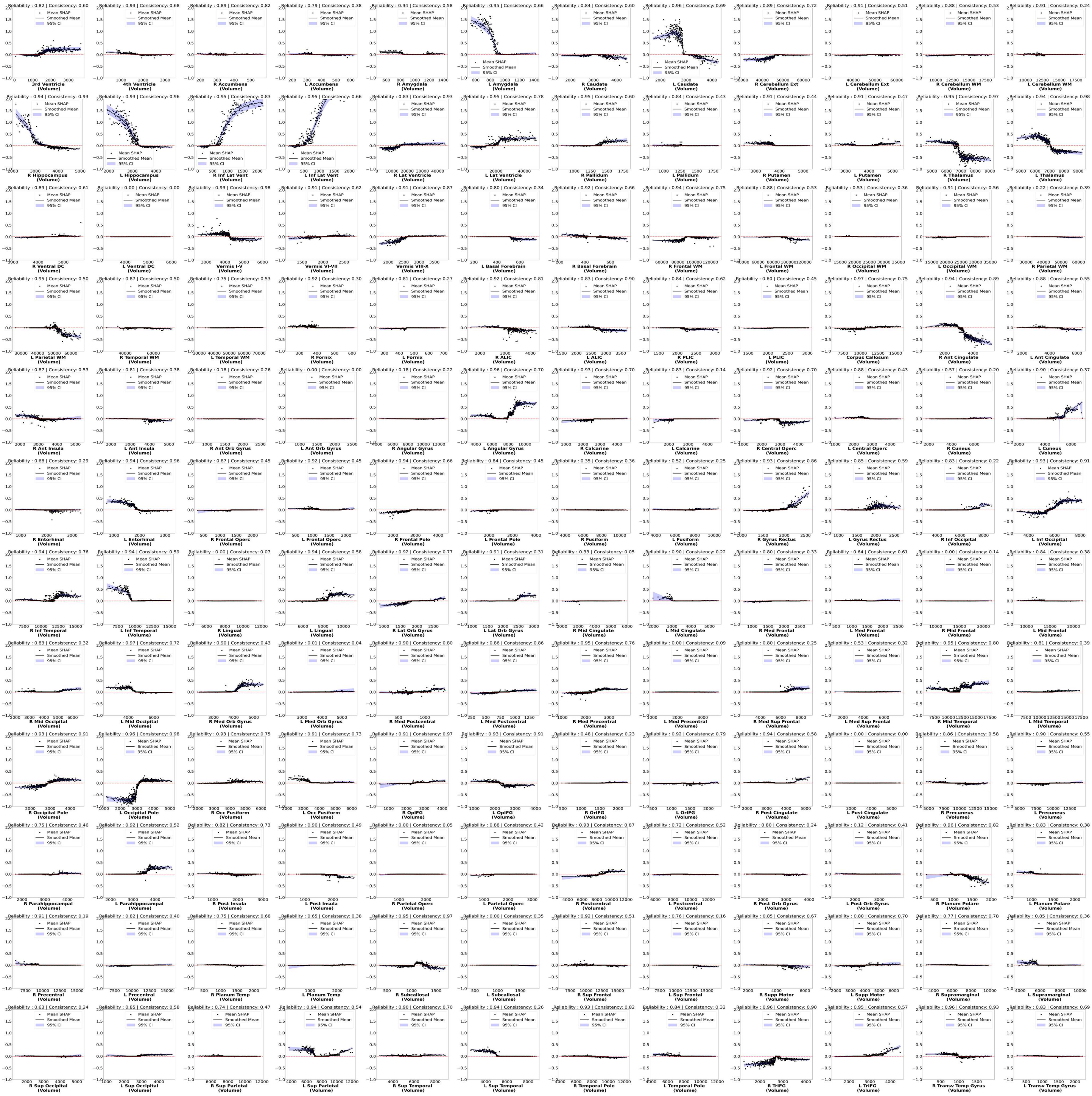

**Figure S6:** (continued) Partial dependence plots for training on NHA.

**(d)** Trained on HISP

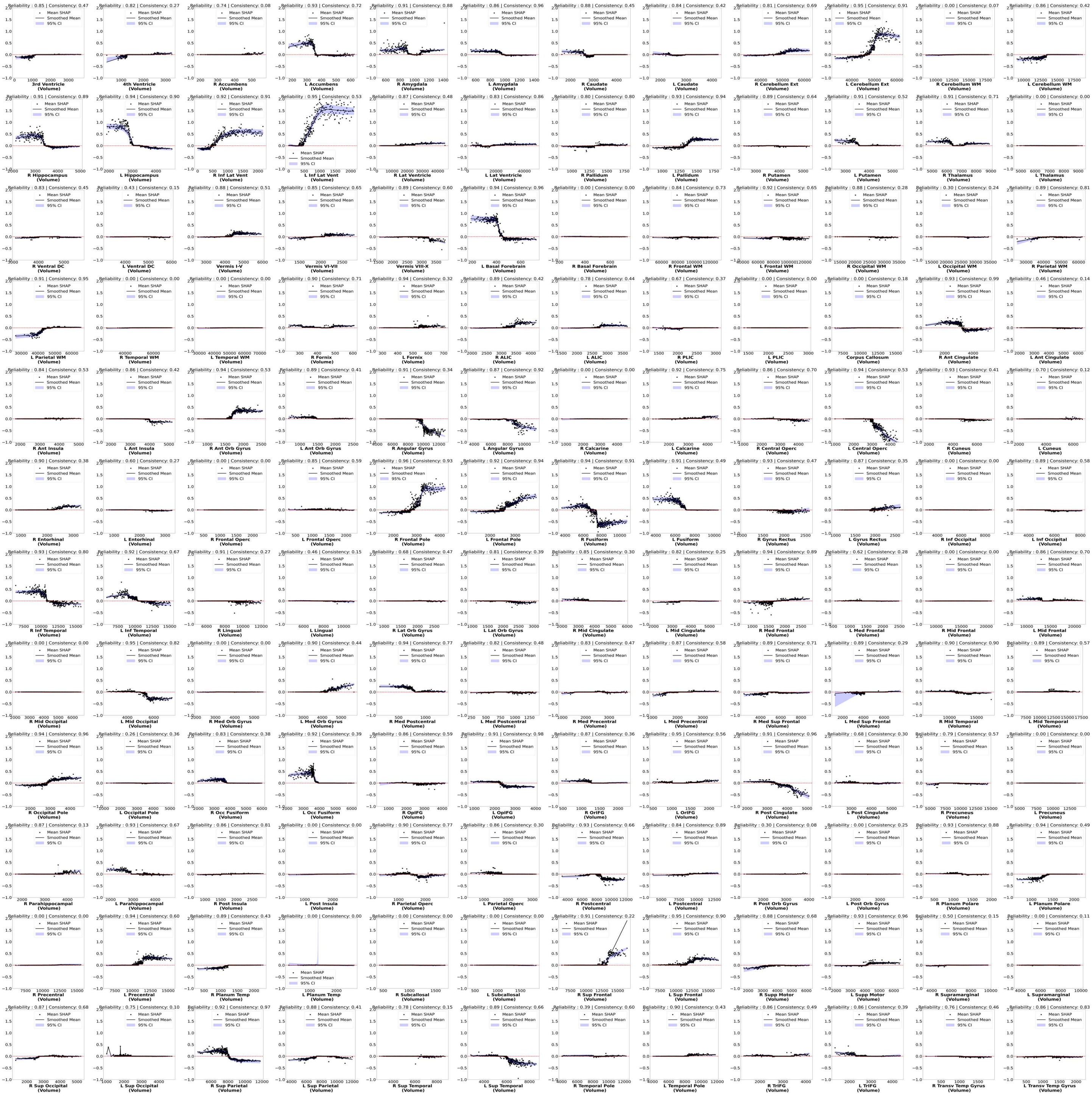

**Figure S6:** (continued) Partial dependence plots for training on HISP.

(**e**) Dictionary of 144 brain regions

| | | | | | | | | | | | |
|---|---|---|---|---|---|---|---|---|---|---|---|
| 3rd Ventricle-VN | 4th Ventricle-VN | R Accumbens-GM | L Accumbens-GM | R Amygdala-GM | L Amygdala-GM | R Caudate-GM | L Caudate-GM | R Cerebellum Ext-GM | L Cerebellum Ext-GM | R Cerebellum WM-WM | L Cerebellum WM-WM |
| R Hippocampus-GM | L Hippocampus-GM | R Inf Lat Vent-VN | L Inf Lat Vent-VN | R Lat Ventricle-VN | L Lat Ventricle-VN | R Pallidum-GM | L Pallidum-GM | R Putamen-GM | L Putamen-GM | R Thalamus-GM | L Thalamus-GM |
| R Ventral DC-WM | L Ventral DC-WM | Vermis I-V-GM | Vermis VI-VII-GM | Vermis VIII-X-GM | L Basal Forebrain-GM | R Basal Forebrain-GM | R Frontal WM-WM | L Frontal WM-WM | R Occipital WM-WM | L Occipital WM-WM | R Parietal WM-WM |
| L Parietal WM-WM | R Temporal WM-WM | L Temporal WM-WM | R Fornix-WM | L Fornix-WM | R ALIC-WM | L ALIC-WM | R PLIC-WM | L PLIC-WM | Corpus Callosum-WM | R Ant Cingulate-GM | L Ant Cingulate-GM |
| R Ant Insula-GM | L Ant Insula-GM | R Ant Orb Gyrus-GM | L Ant Orb Gyrus-GM | R Angular Gyrus-GM | L Angular Gyrus-GM | R Calcarine-GM | L Calcarine-GM | R Central Operc-GM | L Central Operc-GM | R Cuneus-GM | L Cuneus-GM |
| R Entorhinal-GM | L Entorhinal-GM | R Frontal Operc-GM | L Frontal Operc-GM | R Frontal Pole-GM | L Frontal Pole-GM | R Fusiform-GM | L Fusiform-GM | R Gyrus Rectus-GM | L Gyrus Rectus-GM | R Inf Occipital-GM | L Inf Occipital-GM |
| R Inf Temporal-GM | L Inf Temporal-GM | R Lingual-GM | L Lingual-GM | R Lat Orb Gyrus-GM | L Lat Orb Gyrus-GM | R Mid Cingulate-GM | L Mid Cingulate-GM | R Med Frontal-GM | L Med Frontal-GM | R Mid Frontal-GM | L Mid Frontal-GM |
| R Mid Occipital-GM | L Mid Occipital-GM | R Med Orb Gyrus-GM | L Med Orb Gyrus-GM | R Med Postcentral-GM | L Med Postcentral-GM | R Med Precentral-GM | L Med Precentral-GM | R Med Sup Frontal-GM | L Med Sup Frontal-GM | R Mid Temporal-GM | L Mid Temporal-GM |
| R Occipital Pole-GM | L Occipital Pole-GM | R Occ Fusiform-GM | L Occ Fusiform-GM | R OpIFG-GM | L OpIFG-GM | R OrIFG-GM | L OrIFG-GM | R Post Cingulate-GM | L Post Cingulate-GM | R Precuneus-GM | L Precuneus-GM |
| R Parahippocampal-GM | L Parahippocampal-GM | R Post Insula-GM | L Post Insula-GM | R Parietal Operc-GM | L Parietal Operc-GM | R Postcentral-GM | L Postcentral-GM | R Post Orb Gyrus-GM | L Post Orb Gyrus-GM | R Planum Polare-GM | L Planum Polare-GM |
| R Precentral-GM | L Precentral-GM | R Planum Temp-GM | L Planum Temp-GM | R Subcallosal-GM | L Subcallosal-GM | R Sup Frontal-GM | L Sup Frontal-GM | R Supp Motor-GM | L Supp Motor-GM | R Supramarginal-GM | L Supramarginal-GM |
| R Sup Occipital-GM | L Sup Occipital-GM | R Sup Parietal-GM | L Sup Parietal-GM | R Sup Temporal-GM | L Sup Temporal-GM | R Temporal Pole-GM | L Temporal Pole-GM | R TrIFG-GM | L TrIFG-GM | R Transv Temp Gyrus-GM | L Transv Temp Gyrus-GM |

**Figure S6:** (continued) Table of 144 brain regions used in the dementia prediction model. Each cell represents a distinct anatomical region derived from MUSE, encompassing both gray matter (GM), white matter (WM), and ventricular nuclei (VN) labels. Abbreviations include: R/L (Right/Left), Inf (Inferior), Lat (Lateral), Vent (Ventricle), Ant (Anterior), Orb (Orbital), Operc (Operculum), Mid (Middle), Med (Medial), Occ (Occipital), Post (Posterior), Sup (Superior), Supp (Supplementary), ALIC (Anterior Limb of the Internal Capsule), PLIC (Posterior Limb of the Internal Capsule), OpIFG (Opercular Inferior Frontal Gyrus), OrIFG (Orbital Inferior Frontal Gyrus), TrIFG (Triangular Inferior Frontal Gyrus), Temp (Temporal), and Operc (Operculum). This layout corresponds to the order of SHAP values in the above partial dependence plots.

alzheimer's disease. *NeuroImage*, 14(2):298–309.

**Table S2:** VBM studies included in the ALE meta-analysis.

| Reference | Experiment | N | Data Source |
|---|---|---|---|
| Baron et al.[1] | AD < CN | 19 | University of Caen |
| Baxter et al.[2] | AD < CN | 15 | Sun Health Research Institute |
| Berlingeri et al.[3] | AD < CN | 21 | University of Milano-Bicocca |
| Bozzali et al.[4] | AD < CN | 31 | IRCCS Fondazione Santa Lucia |
| Brambati et al.[5] | AD < CN | 10 | McGill Center for Studies in Aging |
| Brenneis et al.[6] | AD < CN | 10 | General Hospital of Linz |
| Canu et al.[7] | EOAD < CN | 18 | IRCCS Centro San Giovanni di Dio FBF |
| Canu et al.[7] | LOAD < CN | 24 | IRCCS Centro San Giovanni di Dio FBF |
| Chen et al.[8] | AD < CN | 15 | University of Pennsylvania |
| Colloby et al.[9] | AD < CN | 47 | Newcastle University |
| Derflinger et al.[10] | AD < CN | 35 | Technische Universität München |
| Farrow et al.[12] | AD < CN | 7 | North Sheffield Research |
| Feldmann et al.[13] | AD < CN | 6 | Local |
| Gee et al.[14] | AD < CN | 12 | University of Pennsylvania |
| Guo et al.[15] | AD < CN | 13 | Xuanwu Hospital |
| Hamalainen et al.[16] | AD < CN | 15 | University of Kuopio |
| Hirao et al.[17] | AD < CN | 61 | National Center Hospital of Neurology and Psychiatry, Tokyo |
| Honea et al.[18] | AD < CN | 60 | University of Kansas Brain Aging Project |
| Hornberger et al.[19] | AD < CN | 15 | FRONTIER database |
| Huang et al.[20] | AD < CN | 50 | Chang Gung Memorial Hospital |
| Ibrahim et al.[21] | AD < CN | 21 | Local |
| Imabayashi et al.[22] | (CT) AD < CN | 5 | Japanese Alzheimer's Disease Neuroimaging Initiative |
| Imabayashi et al.[22] | (MRI) AD < CN | 5 | Japanese Alzheimer's Disease Neuroimaging Initiative |
| Kim et al.[23] | AD < CN | 10 | Chung-Ang University Hospital |
| Kim et al.[23] | AD < CN | 20 | Chung-Ang University Hospital |
| Kim et al.[23] | AD < CN | 31 | Chung-Ang University Hospital |
| Lagarde et al.[24] | AD < CN | 14 | Salpêtrière Hospital |
| Mazere et al.[25] | AD < CN | 8 | University Hospital of Bordeaux |
| Miettinen et al.[26] | AD < CN | 16 | University of Eastern Finland |
| Migliaccio et al.[27] | EOAD Baseline < CN | 15 | University of California San Francisco |
| Migliaccio et al.[27] | LOAD Baseline < CN | 10 | University of California San Francisco |
| Migliaccio et al.[27] | EOAD Progression < CN | 15 | University of California San Francisco |
| Mok et al.[28] | ($CASI_T$) AD < CN | 14 | Shin Kong Wu Ho-Su Memorial Hospital |
| Mok et al.[28] | ($CASI_R$) AD < CN | 10 | Shin Kong Wu Ho-Su Memorial Hospital |
| Polat et al.[29] | AD < CN | 31 | Local |
| Raji et al.[30] | AD < CN | 33 | CHS |
| Rami et al.[31] | AD < CN | 31 | Local |
| Remy et al.[32] | AD < CN | 8 | Local |
| Samuraki et al.[33] | AD < CN | 39 | Kanazawa University Hospital |
| Santos et al.[34] | AD < CN | 34 | Heidelberg University |
| Shiino et al.[35] | AD < CN | 37 | Shiga University of Medical Science |
| Waragai et al.[37] | AD < CN | 15 | Tohoku University |
| Whitwell et al.[38] | Typical AD < CN | 14 | Mayo Clinic |
| Whitwell et al.[38] | Atypical AD < CN | 14 | Mayo Clinic |
| Xie et al.[39] | AD Gray Matter < CN | 13 | Peking University First Hospital |
| Xie et al.[39] | AD White Matter < CN | 13 | Peking University First Hospital |
| Zahn et al.[40] | AD < CN | 10 | University of Freiburg |

EOAD - early onset AD ; LOAD - late onset AD ; $CASI_T$ - Cognitive Abilities Screening Instrument with total scores;
$CASI_R$ - Cognitive Abilities Screening Instrument combined short-term memory and orientation domain scores.
